# Membrane Tension Integrates Physical and Signaling Cues to Gate Cell Fate Transitions

**DOI:** 10.64898/2026.03.04.708749

**Authors:** Gibran Ali, Daniel Gibbard, Elisa Ghelfi, Aaron Olson, Junming Cai, Aidan Balagtas, Isaiah Klein, Emily Maccoux, Yulong Han, Akihiro Miura, Ming Guo, Ian Glass, Munemasa Mori, Douglas G Brownfield

## Abstract

Physical forces shape cell behavior, yet how they integrate with signaling to control fate and disease remains unclear. The alveolar epithelium is patterned by FGF signaling and mechanical stretch, but how these cues specify AT1 and AT2 cells is poorly understood. Here we show that cell membrane tension (CMT) is a conserved regulator of epithelial fate in mouse and human lungs. CMT drops before differentiation and is spatially patterned, defining where bipotent progenitors acquire AT1 or AT2 identity. Lower CMT enhances FGFR2 endocytosis and ERK signaling to drive AT2 differentiation and permits architectural remodeling that enables stretch-mediated YAP/TAZ nuclear entry for AT1 maturation. β-catenin elevates CMT cell-intrinsically independent of its role in canonical WNT transcription, while embedding, osmotic compression, or fibroblast wrapping elevate CMT extrinsically. Combined intrinsic and extrinsic tension traps alveolar epithelial cells in a KRT8⁺ transitional state seen in fibrotic lungs. Membrane tension thus integrates physical and molecular cues linking morphogenesis to fibrosis.

**Graphical Abstract:** 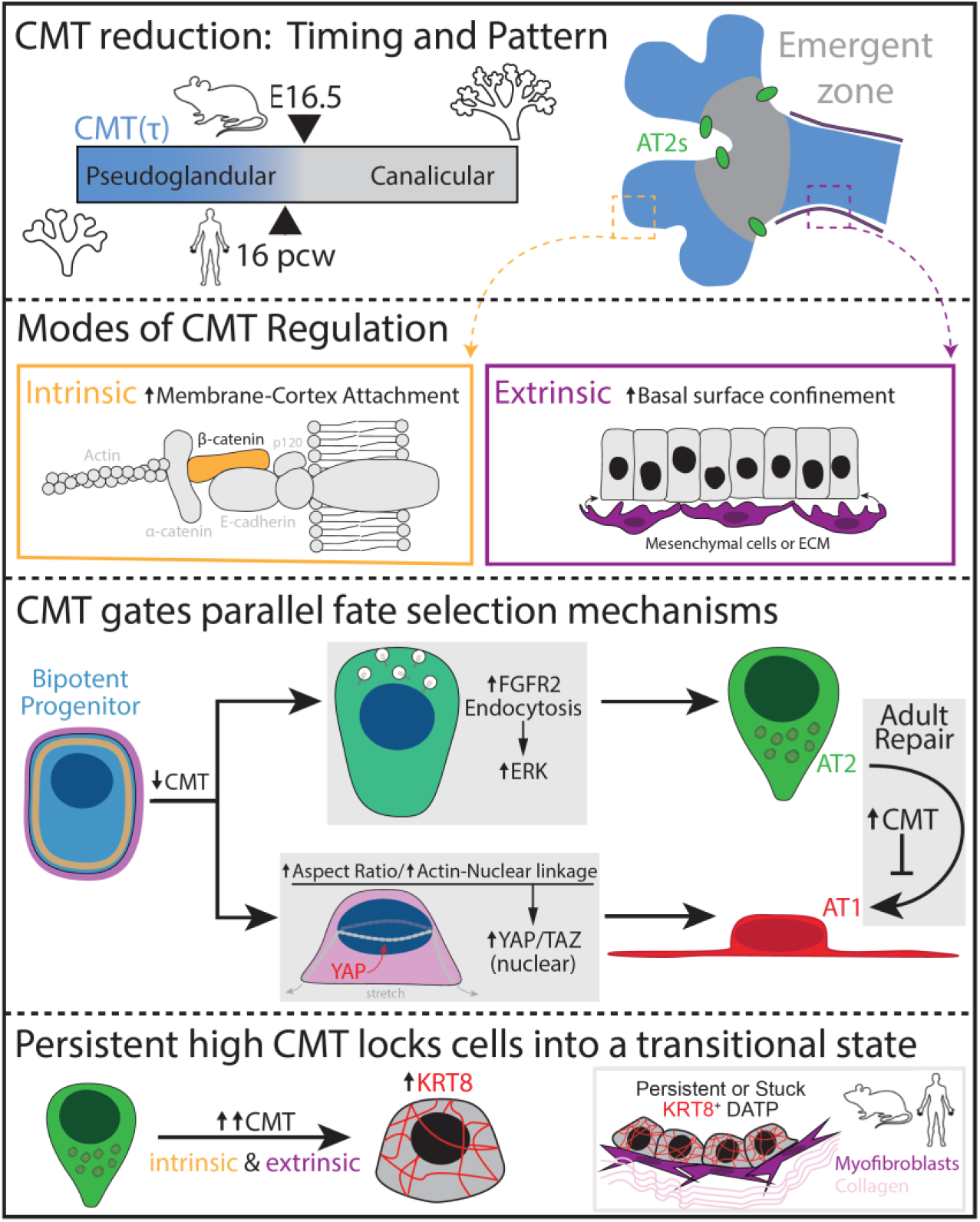

**In Brief:** We identify a conserved drop in cell membrane tension (CMT) that gates epithelial fate transitions across mouse and human lungs. In bipotent progenitors, reduced CMT promotes Alveolar Type 2 (AT2) fate via FGFR2 endocytosis and ERK signaling, as well as Alveolar Type 1 (AT1) fate by enabling architectural remodeling required for YAP/TAZ nuclear entry. We further show that intrinsic β-catenin and extrinsic confinement cues, including mesenchymal contact, converge to elevate CMT, restricting differentiation and—in adult AT2s—inducing a KRT8⁺ transitional state associated with fibrosis.

**Highlights:**

- Cell membrane tension drops before differentiation and is required for AT1 and AT2 fate acquisition.
- Reduced tension enhances FGFR2 endocytosis and ERK signaling to drive AT2 specification.
- Reduced tension permits architectural remodeling and YAP/TAZ nuclear entry required for AT1 maturation.
- Intrinsic β-catenin and extrinsic confinement elevate CMT to restrict differentiation and together strongly induce a KRT8⁺ transitional state associated with fibrosis.

## Introduction

Alveolar epithelial differentiation is a pivotal process during lung development and repair, giving rise to the two morphologically and functionally distinct cell types: the thin, gas-exchanging alveolar type 1 (AT1) cells and the cuboidal, surfactant-producing alveolar type 2 (AT2) cells. Both lineages originate from a bipotent distal progenitor (DP) population, which undergoes lineage segregation into AT2 and AT1 cells to generate mature alveolar epithelium(Desai et al., 2014). Proper coordination of the time and place of progenitor fate specification is essential for establishing functional alveolar architecture, while dysregulation of this process contributes to defective repair and fibrosis(Brownfield et al., 2022; Liberti et al., 2021). Classical studies have defined key molecular regulators of alveolar fate, including FGF signaling, which promotes AT2 differentiation and maintenance(Brownfield et al., 2022), and YAP/TAZ signaling, which drives AT1 specification during late development and persists into adulthood(Gokey et al., 2021; Nantie et al., 2018).

However, molecular cues alone cannot fully account for the temporal delay between the availability of inductive signals (as early as E12) and the emergence of differentiated cells later in development(Treutlein et al., 2014). Indeed, differentiation initiates in spatially patterned “emergent” regions rather than uniformly across the distal epithelium(Sawhney et al., 2025; Zepp et al., 2021). Recent studies point to mechanical forces as critical contributors, especially those extrinsically mediated, such as epithelial–mesenchymal interactions and cyclic respiratory forces(De Belly et al., 2021; Wu et al., 2017). In contrast, intrinsic regulation of the cells’ mechanical properties arises from changes in the composition and connectivity of subcellular components, including the actin cytoskeleton and plasma membrane(Diz-Munoz et al., 2013). Among these intrinsic regulators, cell membrane tension (CMT) has emerged as a critical determinant of stem cell behavior, shown to gate embryonic stem cell (ESC) differentiation(Bergert et al., 2021; De Belly et al., 2021). CMT represents the in-plane resistance of the plasma membrane to deformation and is governed by both lipid composition and degree of membrane-to-cortex attachment (MCA)(Diz-Munoz et al., 2013; Pontes et al., 2017), the latter of which is mediated by membrane–actin linkers such as ERM proteins(Rouven Bruckner et al., 2015).

Elevated CMT has been shown to inhibit endocytosis, restricting fibroblast growth factor receptor (FGFR) internalization and attenuating downstream signaling cascades(De Belly et al., 2021; Riggi et al., 2019; Wu et al., 2017), and can also limit architectural remodeling required for major changes in cell shape(Diz-Munoz et al., 2013). Given the requirement for FGF signaling in alveolar fate control(Brownfield et al., 2022; Liberti et al., 2021; Watson et al., 2022) and the role of cell-shape transitions in AT1 maturation(Li et al., 2018; Shiraishi et al., 2023), CMT represents a plausible integrator of mechanical state and signaling output. In disease contexts, disruptions in the balance of cellular and tissue-level mechanics can lock alveolar epithelium in aberrant transitional states, exemplified by a KRT8⁺ transitional state (often described as a PATS/DATP-like state) observed during injury and reported to persist in idiopathic pulmonary fibrosis (IPF)(Choi et al., 2020; Jiang et al., 2020; Strunz et al., 2020; Wang et al., 2023). The enrichment of KRT8⁺ transitional cells in fibrotic, mechanically rigid regions suggests that high-tension environments may reinforce lineage arrest(Strunz et al., 2020; Wang et al., 2023). Given that CMT functions as a mechanical checkpoint for cytoskeletal remodeling, receptor trafficking, and lineage transitions(Diz-Munoz et al., 2013), we sought to investigate how intrinsic and extrinsic regulation of CMT governs alveolar differentiation during development and how its dysregulation contributes to disease-associated states.

Here, we demonstrate that CMT in distal lung progenitors decreases prior to, and is required for, alveolar epithelial differentiation. Reduced CMT facilitates FGFR2 endocytosis and ERK activation to promote AT2 fate, and enables the architectural remodeling associated with YAP/TAZ nuclear entry and AT1 specification. We demonstrate that CMT is regulated cell-intrinsically via β-catenin, independently of its role in canonical downstream WNT signaling. Further, CMT is regulated in a cell-extrinsic manner through matrix context and epithelial-mesenchymal interactions, both of which impede CMT reduction and alveolar epithelial differentiation. Manipulating CMT through intrinsic and extrinsic inputs modulates AT1 differentiation in human and mouse AT2 cells. Finally, combined intrinsic and extrinsic CMT increase induces KRT8 expression, marking a “stuck” transitional state commonly associated with pulmonary fibrosis(Choi et al., 2020; Jiang et al., 2020; Strunz et al., 2020; Wang et al., 2023). Together, our findings establish CMT as a critical, previously unrecognized regulator of alveolar epithelial fate, revealing a conserved mechano-signaling mechanism that governs both developmental lineage transitions and regenerative outcomes in the adult lung.

## Results

### A conserved and patterned CMT drop precedes and gates alveolar differentiation

To define when CMT changes during alveolar lineage specification, we labeled live embryonic mouse lung slices (E15.5–E17.5) with the mechanosensitive membrane probe Flipper-TR(Colom et al., 2018) and quantified fluorescence lifetime by FLIM (Figure 1A). Across development, CMT progressively decreased in the distal epithelium, with the largest reduction observed between E15.5 and E17.5 (Figure 1B). To validate this tissue-level readout with an orthogonal mechanical measurement, we isolated DPs and nAT2s by MACS and quantified CMT by optical-trap tether rupture (Figure 1C). Early DPs (E15.5) exhibited the highest CMT, which declined in late-stage DPs (E16.5) and remained low in nAT2s (Figure 1D), indicating that the CMT drop precedes differentiation.

**Figure 1:**
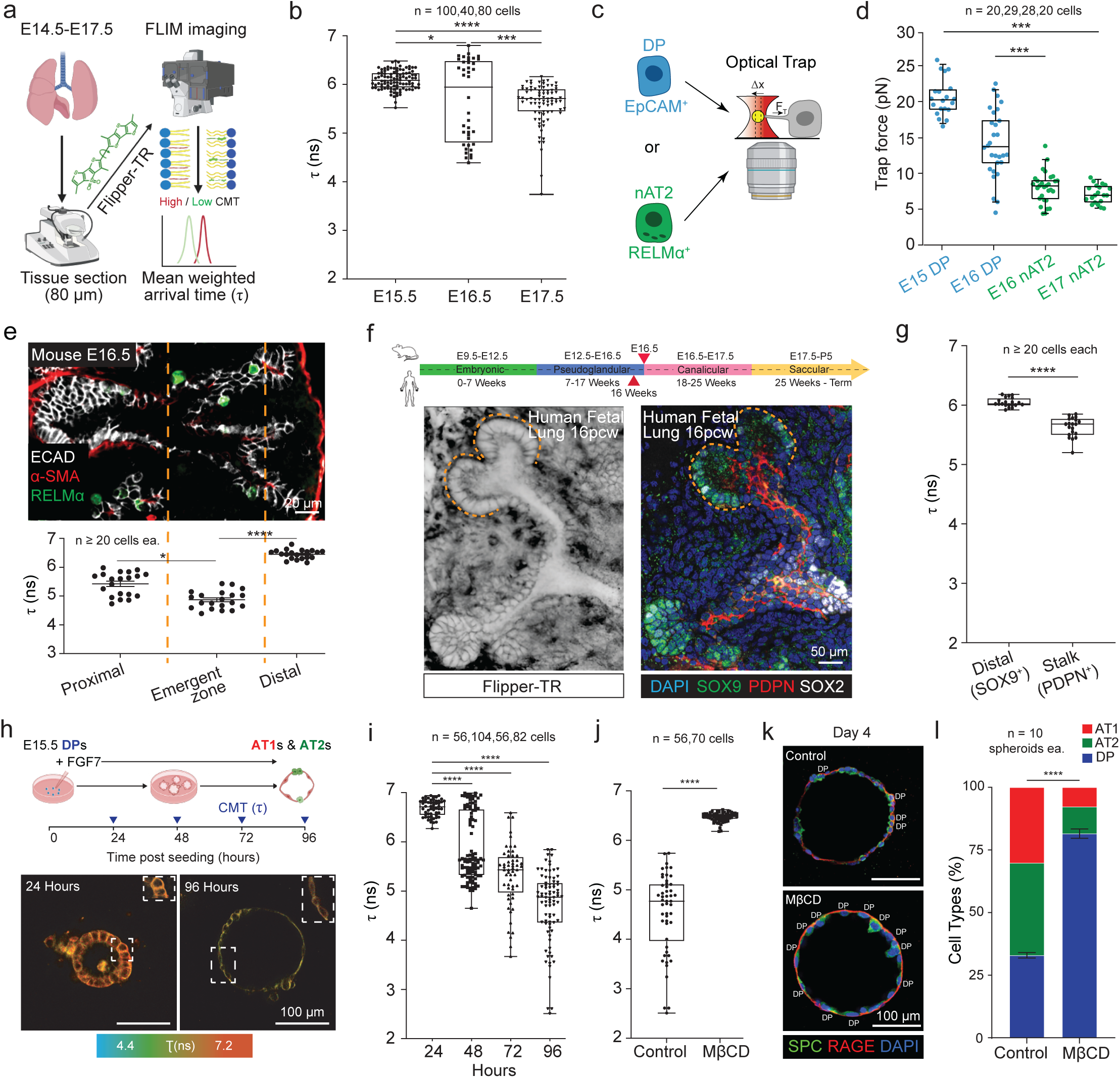
Cell membrane tension reduces prior to and is required for alveolar epithelial differentiation. (A) Approach to measure in situ CMT changes in unfixed living fetal lung with Flipper-TR dye via fluorescence lifetime to determine mean weighted arrival time (or τ). (B) CMT measurements of the mouse distal lung epithelium at developmental timepoints prior to (E15.5) and during (E16.5, E17.5) differentiation. Data represents ± SD of mean combined from 2 independent experiments (p-values *0.0162, **0.0027, and ****<0.0001 from one-way ANOVA with multiple comparisons test). (C) Approach of alveolar cell purification by MACS for CMT measurement via optical trap. (D) Trap force of undifferentiated DPs (blue) prior to (E15) and during (E16) differentiation as well as newly formed nAT2s (green; E16, 17). Data shown as mean ± SD (p-value ***p<0.001 determined using one-way ANOVA). (E) Upper, immunostain of the E16.5 mouse lung epithelium depicting the proximal (basal α-SMA+ ASMs), distal, and recently described emergent zone where RELMα+ AT2s emerge. Lower, in situ CMT measurements at these sites. Data are presented as mean ± SD (p-values *p = 0.0283 and ****p<0.0001 calculated using one-way ANOVA). (F) Upper, matched timelines for mouse and human lung development, red arrows indicate timepoint for human CMT measurement and its closest matched mouse timepoint (E16.5). Left, matched immunostain of 16 pcw fetal lung epithelium following live Flipper-TR imaging, wherein undifferentiated SOX9+ DPs are detected at the tips and SOX9-SOX2-differentiating alveolar epithelial cells in the stalk. (G) CMT measures from tip and stalk regions retrospectively identified for SOX9/SOX2 by matched immunostains. Data are mean ± SD (p-value p < 0.0001 was determined using the Student’s t-test). (H) Upper, experimental timeline of alveolosphere differentiation assay and CMT measurements. Lower, representative fluorescent lifetime images of spheroids after 24 hours and 96 hours in culture. (I) CMT of alveolospheres at 24, 48, 72, and 96 hours in culture. Data are shown as means ± SDs from 4 independent experiments (p-values ****p < 0.0001 determined via two-way ANOVA). (J) CMT was significantly increased in spheroids following treatment with MβCD (1mM). Data are mean ± SD from 3 independent experiments (****p <0.0001 was calculated using a Mann-Whitney test). (K) Immunostains of alveolospheres in either control differentiation conditions alone (50 ng/ml FGF7) or with MβCD. Staining allows for detection of AT1s (RAGE+), AT2s (SPC+), and DPs (RAGE+ SPC+). Scale bars indicate 100µm. (L) Quantification of (K) illustrating a reduction in AT1 and AT2 differentiation, assessed by percentage of DPs per spheroid with three experimental replicates. Statistical significance was determined using a two-tailed Mann-Whitney test, yielding a p-value of <0.0001.

We next asked whether this reduction is spatially patterned within the developing lung. At E16.5, CMT exhibited a reproducible proximal–distal pattern: regions corresponding to the emergent zones where alveolar differentiation initiates displayed the lowest CMT, more proximal regions were intermediate, and distal tips retained the highest CMT (Figure 1E). We then tested whether this spatial pattern is conserved in human development. In human fetal lung at 16 post-conception weeks (pcw), CMT similarly varied along the distal–proximal axis, with lower CMT in PDPN⁺ stalk regions and higher CMT at SOX9⁺ distal tips (Figure 1F–G), consistent with conserved spatial regulation of membrane mechanics.

Finally, we asked whether the developmental CMT drop is required for alveolar epithelial differentiation. In an organotypic assay, E15.5 DPs stimulated with FGF7 reduced CMT within 48 hours in culture (Figure 1H), preceding the first detection of AT1 or AT2 markers in this system (Figure S1A–C). Preventing this drop by elevating CMT with methyl-β-cyclodextrin (MβCD, 1 mM) maintained higher CMT and markedly impaired differentiation into both AT1 and AT2 fates following FGF7 stimulation (Figure 1J–L). In parallel, human fetal distal epithelial cells (16 pcw) robustly differentiated into SPC⁺ AT2 cells in AT2 differentiation medium, but this transition was suppressed when CMT was increased with polyethylene glycol (1.5% PEG) (Figure S1E–F). Together, these data show that CMT decreases in a conserved, spatially patterned manner prior to alveolar fate acquisition and that maintaining high CMT inhibits alveolar epithelial differentiation.

### CMT reduction enables endocytic FGFR2–ERK signaling required for AT2 specification

Having established that lowering CMT promotes alveolar epithelial differentiation, we asked which downstream processes are being gated. Because elevated CMT can suppress endocytosis during differentiation(De Belly et al., 2021), we first quantified endocytic activity in DPs across late embryonic development. EEA1^+^ early endosomes increased markedly between E15.5 and E17.5 in both DPs and nAT2s (Figure 2A–B), coincident with the developmental reduction in CMT. In culture, nAT2s similarly exhibited higher uptake of a pH-sensitive dextran tracer than DPs (Figure S2A–B). Functionally, elevating CMT with MβCD (1 mM) reduced EEA1 puncta in differentiating DPs (Figure 2C–D), while inhibiting dynamin-dependent endocytosis with Dynasore (10 μM) strongly impaired acquisition of both AT1 and AT2 fates (Figure 2E–F). Together, these data indicate that endocytosis increases as CMT decreases and is required for alveolar epithelial differentiation.

**Figure 2:**
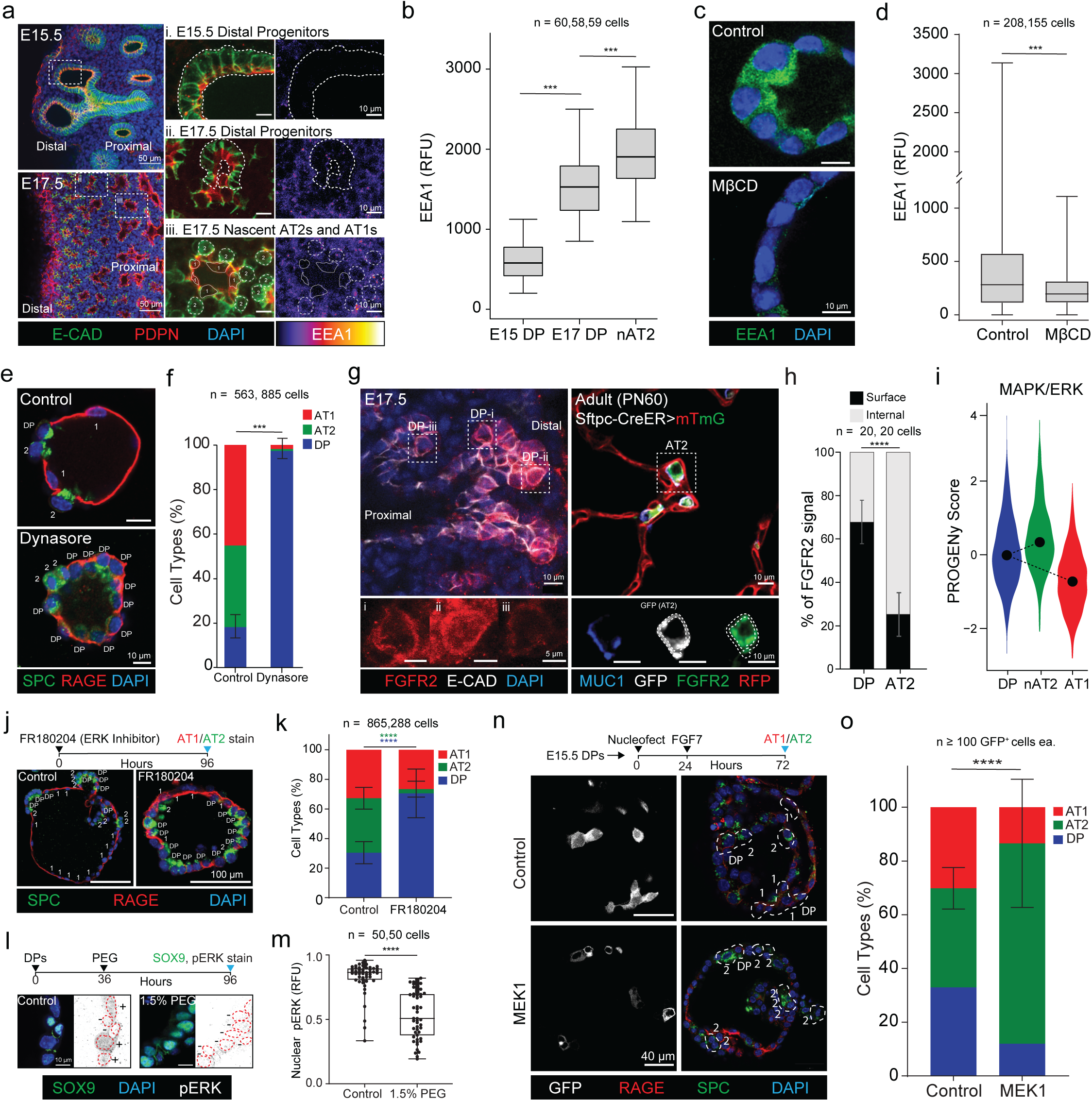
CMT reduction drives FGFR2 endocytosis, ERK signaling, and AT2 fate selection. (A) Immunostains of E15.5 and E17.5 mouse distal lung epithelium wherein early endosomes (EEA1) are detected in DPs and nAT2s. AT1s (1) and AT2s (2) are identifiable at E17.5 by morphology and podoplanin (PDPN, red). Scale bars correspond to 50µm and 10µm, respectively. (B) Quantification of EEA1 in late DPs as well as nAT2s. Error bars show 95% confidence intervals; n ≥ 58 cells (p-values ***p < 0.001 calculated via one-way ANOVA). (C) Immunostain of EEA1 in alveolospheres 4 days in differentiation media treated with DMSO (Control) or raised CMT (MβCD, 1mM), scale bar represents 10µm. (D) Quantification of c showing a reduction of EEA1 when CMT is raised. Expressed as mean ± SD from three independent experiments (Control: n = 208 cells; MβCD: n = 155 cells). Statistical significance was assessed using a two-tailed Mann-Whitney test, with results indicating ***p < 0.001. (E) Immunostains of alveolospheres at day 4 following treatment with DMSO (Control) or 10 µM Dynasore for markers of AT1 (RAGE, red) and AT2 (SPC, green) fate. Scale bars indicate 10µm. (F) Quantification of e detects a significant reduction in differentiation with Dynasore treatment. Data are presented as mean ± SD (p-value ***p < 0.001 determined by two-tailed Mann-Whitney test). (G) FGFR2 immunostains of E17.5 (left) and adult (right) mouse lungs, wherein the cell surface can be detected by either E-Cadherin or membrane GFP. Lower breakouts highlight FGFR2 cell surface enrichment at E17.5 (i-iii) that transitions to intracellular (lower right, surface depicted by dashes) in adult AT2s. (H) Quantification of g finds a significant shift in localization over time. Data are mean ± SD; n = 20 cells per group (p < 0.0001 determined using a two-tailed Mann-Whitney test). (I) PROGENy score of ERK pathway activity in mouse DPs, nAT2s and AT1s sampled in a prior scRNAseq time course study. (J) Alveolospheres at day 4 under differentiation conditions treated with DMSO (Control) or ERK inhibitor (FR180204, 10µM) stained for AT markers, scale bar 100μm. (K) Quantification of j, demonstrating a significant reduction in AT2 differentiation. Data shown as mean ± SD; n = 10 spheroids per condition conducted in triplicate (p-values p< 0.0001 were determined using a 2-way ANOVA). (L) Differentiating alveolospheres in Control or raised CMT conditions (1.5% PEG) stained for DP marker (SOX9) and phosphorylated ERK (nuclei in red, pERK positivity/negativity indicated as +/-), scale 10μm. (M) Quantification of l nuclear pERK intensity. Data expressed as mean ± SD; n = 50 cells per group (p-value p < 0.0001 determined by two-tailed Mann-Whitney test). (N) Alveolospheres wherein either GFP or MEK1 was mosaically expressed then immunostained for AT markers, scale bar indicates 100μm. (O) Quantification of GFP+ cells from (N) finds a significant upregulation of AT2 fate in MEK1 expressing cells versus control. Data presented as mean ± SD. p-value p < 0.0001 was determined by two-tailed Mann-Whitney test.

Endocytosis can also tune growth factor signaling by controlling receptor trafficking(Sorkin and von Zastrow, 2009). Because FGFR2 signaling is a central determinant of AT2 identity(Brownfield et al., 2022) and ERK activation can depend on receptor internalization(De Belly et al., 2021), we next tested whether the CMT-dependent increase in endocytosis enables an FGFR2–ERK program during AT2 specification. Immunostaining revealed a shift in FGFR2 localization from predominantly surface-associated in E17.5 DPs to a punctate intracellular pattern in mature adult AT2s (Figure 2G–H), consistent with increased receptor internalization during AT2 maturation. In parallel, analysis of a published mouse lung developmental scRNA-seq time course using PROGENy/decoupleR inferred a selective increase in MAPK/ERK pathway activity as cells transitioned into the AT2 state (Figure 2I; Figure S2F). Consistent with a functional requirement, ERK inhibition (FR180204, 10 μM) significantly reduced AT2 differentiation, with a stronger effect than inhibitors of other FGF-associated downstream pathways (Figure 2J; Figure S2C–E). Importantly, elevating CMT (1.5% PEG) reduced pERK levels in differentiating DPs (Figure 2L–M), indicating that raised CMT suppresses ERK activation. Conversely, enforcing downstream ERK pathway activity by nucleofection of MEK1-GFP increased AT2 differentiation among GFP^+^ cells relative to vector controls (Figure 2N–O). Collectively, these results support a model in which developmental reduction in CMT relieves an endocytic constraint, enabling FGFR2–ERK signaling that is required—and when activated downstream, sufficient—to promote AT2 specification.

### CMT reduction enables YAP nuclear entry and AT1 specification

YAP/TAZ-mediated mechanotransduction is a key driver of AT1 maturation, which is accompanied by extensive architectural remodeling of the cytoskeleton and nucleus. Consistent with this, mechanotransduction-associated components are enriched along the DP→AT1 trajectory (e.g., integrin/cytoskeletal and nuclear-coupling factors; Figure S3A). To directly assess whether AT1 emergence involves nuclear remodeling in our differentiation system, we quantified nuclear shape and actin association across DP, AT2, and AT1 states. Marker-defined AT1 cells exhibited a marked increase in nuclear aspect ratio and increased nuclear–actin overlap relative to DPs and AT2 cells (Figure 3A–C).

**Figure 3:**
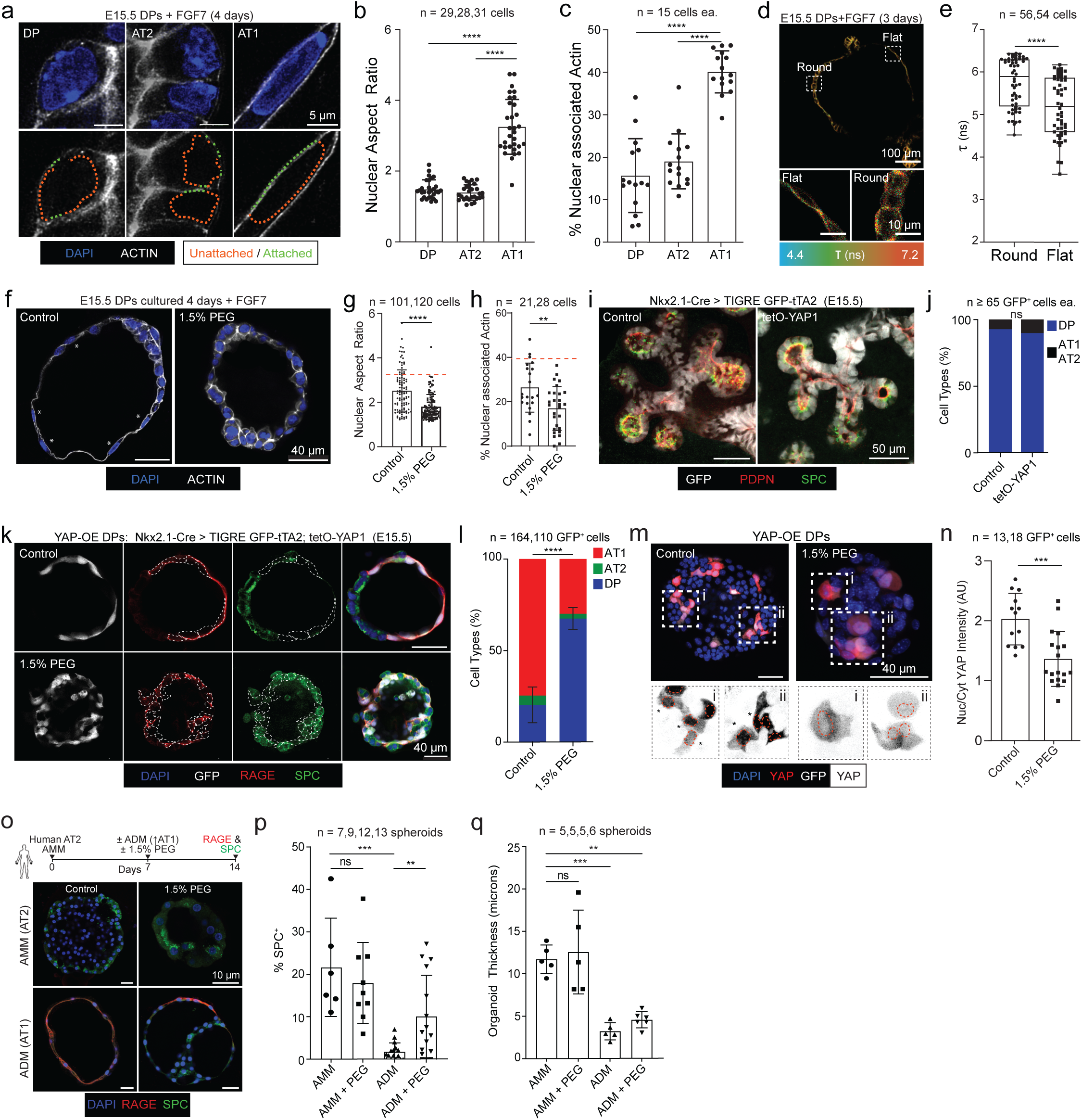
Raised CMT suppresses architectural remodeling of the nucleus necessary for YAP internalization and AT1 differentiation. (A) Representative DP, AT2 and AT1 nuclear morphologies. Dotted green line indicates DAPI and Phalloidin overlap, orange no overlap (scale bar, 5μm). (B) Quantification of nuclear aspect ratio from a. Data presented as mean ± SD, p-values **p < 0.01, ****p < 0.0001 were calculated using one-way ANOVA. (C) Quantification of nuclear-actin overlap in a. Data presented as mean ± SD, p-values ****p < 0.0001, ***p < 0.001 calculated via one-way ANOVA. (D) Flipper-TR stain of alveolospheres after 3 days in differentiation media wherein both flat (likely AT1) and round cells (likely AT2) can be detected (scale bars, 100μm above and 10μm lower). (E) Quantification of d detects a further CMT reduction in flattened (likely AT1) cells (p-value ****<0.0001 calculated by Mann-Whitney test). (F) Alveolospheres 4 days in differentiation media under either Control (DMSO) or raised CMT (1.5% PEG) conditions stained with DAPI and Phalloidin (scale bar, 40μm). (G) Quantification of nuclear aspect ratio from e with average AT1 value from b in dashed red, each sampled in experimental triplicate (p-value ****p < 0.0001 was calculated by Mann-Whitney test). (H) Quantification of nuclear-actin attachment from f with average AT1 value from c in dashed red: each sampled in experimental triplicate (p-value calculated via Mann-Whitney test. (I) Transgenic E15.5 mouse lungs wherein the distal epithelium was either GFP labeled alone (*Tg^Nkx2.1-Cre^*; *Tigre^GFP-tTA2^*, Control) or also constitutively express YAP1 (*Tg^Nkx2.1-Cre^*; *Tigre^GFP-tTA2^*; *Rosa26^tetO-S112A-YAP1^*, tetO-YAP1) stained for restrictive fate markers SPC and PDPN (scale bar, 50µm). (J) Quantification of DPs (SPC^+^ PDPN^+^) and differentiated AT1s or AT2s (either SPC+ or PDPN+) from i (p-value ns, Student’s t-test). (K) E15.5 DPs isolated from tetO-YAP1 lungs under differentiation media for four days in either control or raised CMT (1.5% PEG) conditions, then stained for SPC and RAGE (scale bar, 40µm). Mosaic GFP^+^ cells from Control or YAP expressing conditions are indicated in dashed lines. (L) Quantification GFP^+^ cell proportion from k (p-value ****p<0.0001, Student’s t-test). (M) YAP immunostains from experiment in k with example GFP+ cells indicated (i, ii), nuclei are in dashed red and asterisks indicate enriched nuclear YAP (scale bar, 40µm). (N) Quantification of YAP (cell proportion with enriched nuclear YAP) from m (p-value calculated using a Mann-Whitney test). (O) Above; Experiment timeline for human AT2s treated with Alveolar Maintenance Media (AMM, reinforces AT2 differentiation) and Alveolar Differentiation Media (ADM, promotes AT2-to-AT1 differentiation) with or without raised CMT (1.5% PEG). (Lower) alveolospheres from stated conditions stained for SPC and RAGE. (P) Quantification of SPC positivity of spheroids in (O). (Q) Quantification of average organoid thickness from o. p-values ****p < 0.0001, ***p < 0.001, **p < 0.01 calculated via two-way ANOVA.

We next asked whether changes in CMT precede these architectural transitions. At day 3 of differentiation—when both flattened and rounded epithelial morphologies are present—flattened cells exhibited significantly lower Flipper-TR lifetime (lower CMT) than neighboring rounded cells (Figure 3D–E), indicating that CMT reduction accompanies the onset of AT1-like morphological remodeling. Conversely, elevating CMT with PEG prevented nuclear elongation and reduced nuclear–actin association during differentiation (Figure 3F–H), demonstrating that reduced membrane tension is required for the cytoskeletal–nuclear remodeling characteristic of AT1 maturation.

Because YAP activity depends on mechanical context and nuclear access(Lopez et al., 2011; Zhang et al., 2021), we tested whether high CMT suppresses YAP nuclear accumulation during AT1 specification. We first asked whether enforced YAP expression is sufficient to induce AT1 fate at E15.5, a developmental stage at which distal progenitors retain elevated CMT. Using a tetO-YAP1 allele(Lim et al., 2023) driven in Nkx2.1-lineage^+^ distal epithelium, YAP expression did not alter epithelial morphology or the prevalence of transitional SPC⁺PDPN⁺ DPs at E15.5 (Figure 3I–J), indicating that YAP expression alone is insufficient to trigger AT1 fate under high-CMT conditions. We next tested CMT regulation of AT1 fate and YAP in vitro. E15.5 DPs isolated from tetO-YAP1 embryos differentiated robustly toward RAGE⁺ AT1 cells under control conditions; however, elevating CMT with PEG markedly reduced AT1 differentiation and increased retention of DP-like states (Figure 3K–L). Importantly, despite enforced YAP expression, high CMT significantly reduced YAP nuclear accumulation as quantified by nuclear-to-cytoplasmic YAP intensity (Figure 3M–N). We observed similar behavior when YAP protein levels were increased pharmacologically via LATS inhibition with a small molecule (TRULI): elevated CMT suppressed both AT1 differentiation and YAP nuclear accumulation (Figure S3B–E).

Finally, we asked whether CMT similarly constrains AT1 maturation in adult human tissue. In an adult human AT2-to-AT1 differentiation assay, alveolar differentiation medium (ADM) reduced SPC positivity and decreased organoid thickness relative to maintenance conditions, consistent with AT1 maturation. Elevating CMT with PEG partially reversed these ADM-driven changes, increasing SPC positivity and organoid thickness (Figure 3O–Q). Together, these results identify CMT reduction as a mechanical checkpoint that permits cytoskeletal–nuclear remodeling and thereby enables YAP nuclear accumulation required for AT1 fate acquisition.

### β-catenin maintains progenitor status by elevating CMT independent of WNT transcription

Having found that a reduction in CMT is required for both AT1 and AT2 maturation, we next asked how it is maintained at high levels in early DPs. Because CMT reflects the balance between actomyosin contractility and membrane–cortex attachment(Diz-Munoz et al., 2013), we first tested whether increasing membrane–cortex coupling is sufficient to raise CMT and impede differentiation. We mosaically expressed a synthetic membrane–cortex linker (iMC-linker(Bergert et al., 2021)) that mechanically couples the plasma membrane to cortical actin and quantified CMT specifically in labeled cells. iMC-linker expression significantly increased Flipper-TR lifetime relative to membrane-targeted mCherry controls (Figure 4A), demonstrating that enhanced membrane–cortex attachment is sufficient to elevate CMT in a cell-intrinsic manner. Consistent with a mechanical “gate” on fate acquisition, iMC-linker–expressing cells were strongly biased toward a DP-like state and showed reduced differentiation into SPC⁺ AT2 and RAGE⁺ AT1 cells (Figure 4B–C).

**Figure 4:**
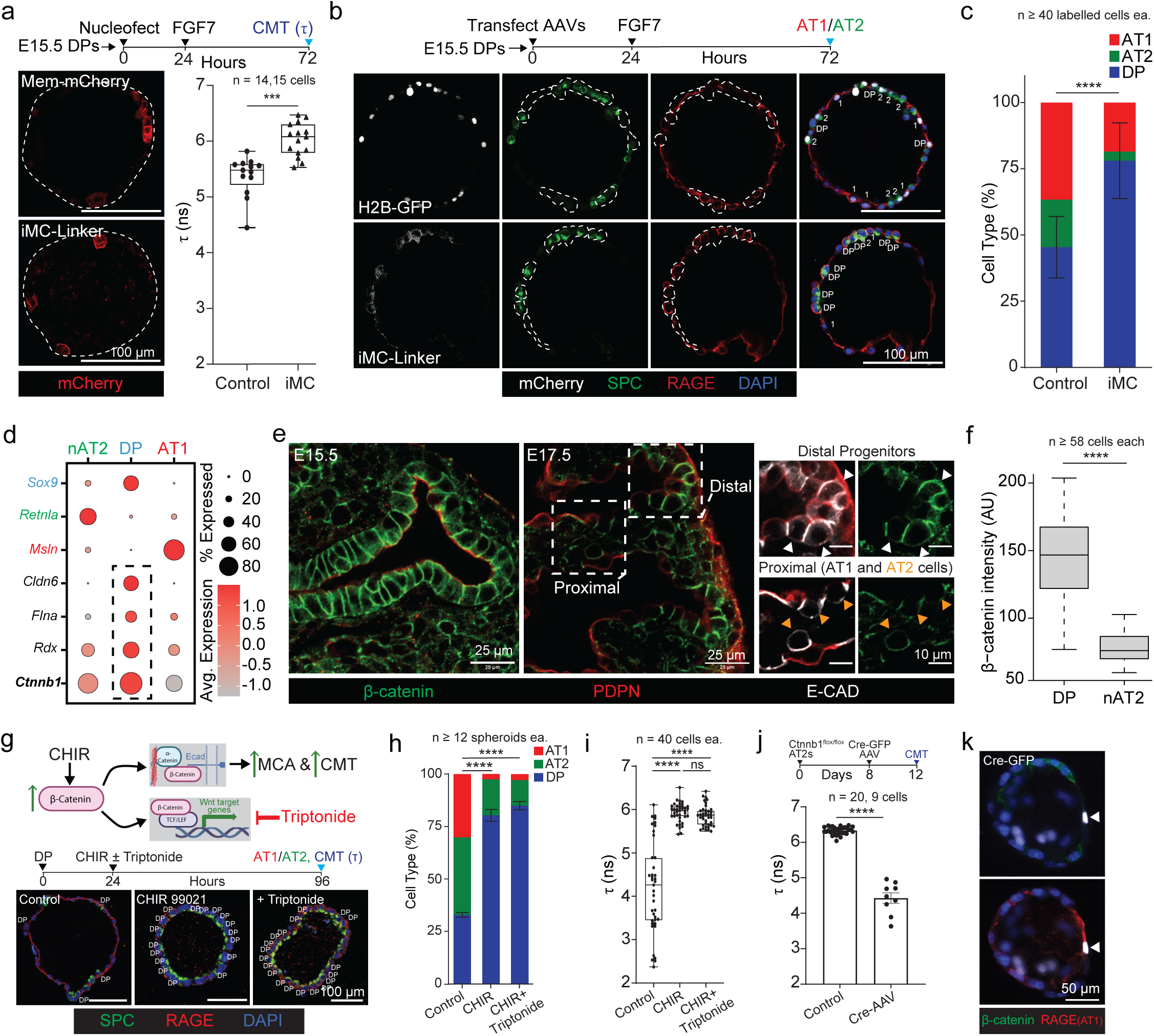
β-Catenin is an intrinsic regulator of CMT that blocks fate transitions independent of downstream WNT signaling. (A) (Left) Alveolospheres mosaically expressing either control (Mem-mCherry) or cell intrinsic CMT raising construct (iMC-Linker) three days in culture (scale bar, 50μm). (Right) CMT quantification of mosaically labeled mCherry^+^ cells. Data shown as mean ± SD (p-value ***p < 0.001, two-tailed Mann-Whitney test). (B) Alveolospheres from control or iMC-Linker conditions stained for SPC, RAGE, after three days in differentiation media (scale bars, 100µm). (C) Quantification of cell types from b of Control (H2B-GFP) and iMC-Linker (iMC) conditions. Cells sampled per condition in experimental duplicate (p-value ****p < 0.0001, two-tailed Mann-Whitney test). (D) Dot plot from a time course scRNAseq dataset sampling mouse lung development wherein marker genes for nAT2 (green), DP (blue), and AT1 (red) are colored and genes encoding proteins that putatively regulate CMT in black, dashed box indicates upregulation in DPs (gene encoding β-catenin in bold). (E) Representative immunostains of β-catenin prior to (E15.5) and during (E17.5) alveolar epithelial differentiation in the mouse lung. Costaining for E-Cadherin and Podoplanin allow for identification (along with timepoint and anatomical position) of DPs (white arrowheads), nAT2s (orange arrowheads), and AT1s (scale bars, 25µm and 10µm, respectively). (F) Quantification of β-catenin intensity from E17.5 DPs and nAT2s from e. Data shown as mean ± SD, (p-value ****p < 0.0001, two-tailed Mann-Whitney test). (G) (Top) Schematic of experiment wherein β-catenin is upregulated via CHIR treatment (CHIR99021, 3µM) but its downstream role in transcriptional regulation of WNT genes suppressed via small molecule (Triptonide, 100nM), leaving intact its membrane-to-cortical actin (MCA) linkage critical for CMT. (Bottom) Immunostains of alveolospheres under the above conditions stained for AT fate markers SPC and RAGE after 4 days in culture (scale bar, 100μm). (H) Quantification of cell type proportion from G per condition. Spheroids sampled per condition in experimental triplicate (p-values ****p < 0.0001, two-way ANOVA). (I) Quantification of CMT from experiment in g (p-values ****p < 0.0001, ns p > 0.05, two-way ANOVA). (J) Upper; Experiment timeline of isolated transgenic mouse β-catenin^fl/fl^ AT2 cells treated with Cre-GFP AAV to label AT2s and deplete β-Catenin to assess its necessity for raised CMT and fate maintenance in the presence of CHIR. (Bottom) CMT quantification finds a significant reduction in AT2s wherein β-catenin is removed. Data shown as mean ± SD (p-values ****p < 0.0001, two-tailed Mann-Whitney test). (K) Representative GFP^+^ β-catenin^fl/fl^ AT2 cell (arrowhead) that lacks β-catenin and has a flattened and RAGE+ AT1 phenotype (scale bar, 50μm).

To identify endogenous regulators that sustain high CMT in early DPs, we analyzed a developmental scRNA-seq time course and found several candidate membrane/cortex-associated factor genes enriched in DPs, including *Cldn6*, *Rdx*, and *Ctnnb1* (β-catenin) (Figure 4D). Given the known roles of β-catenin in both canonical WNT transcription and adherens junction organization(Valenta et al., 2012), we asked whether β-catenin abundance correlates with the high-CMT progenitor state in vivo. Immunostaining revealed high β-catenin levels in E15.5 distal epithelium and a reduction in β-catenin as cells transition toward nAT2 identity at later stages (Figure 4E–F). We next tested whether elevating β-catenin is sufficient to maintain high CMT and block differentiation, and whether this requires canonical WNT/TCF transcriptional output. Stabilizing β-catenin with the GSK-3β inhibitor CHIR99021 increased CMT and strongly inhibited AT1/AT2 differentiation. Importantly, co-treatment with Triptonide—which suppresses β-catenin–dependent WNT/TCF transcriptional output—did not reduce the CHIR-induced increase in CMT or rescue differentiation (Figure 4G–I). Taken together, this indicates that β-catenin can maintain a high-CMT state through a transcription-independent mechanism consistent with its junctional role.

Finally, we tested whether β-catenin is necessary to sustain elevated CMT. In cultured *Ctnnb1^flox/flox^* AT2 cells maintained under CHIR conditions, mosaic deletion of β-catenin by Cre-GFP AAV significantly reduced CMT (Figure 4J) and was associated with cell flattening and acquisition of an AT1-like RAGE⁺ phenotype in β-catenin–deleted GFP⁺ cells (Figure 4K). Collectively, these results identify β-catenin as an intrinsic regulator of membrane mechanics that sustains high CMT to restrict epithelial fate transitions, largely independent of canonical WNT transcriptional output.

### Extrinsic confinement and mesenchymal contact elevate CMT to restrict differentiation

Having shown that elevated CMT is sufficient to block AT1 and AT2 maturation, we next asked whether extrinsic physical cues modulate CMT to influence progenitor fate. In prior work we observed that DP organoids differentiate more robustly when cultured on top of Matrigel(Brownfield et al., 2022; Sawhney et al., 2025) rather than embedded, suggesting that physical context may impact differentiation. We therefore compared CMT and fate outcomes between these culture geometries. DPs cultured on top of Matrigel exhibited a pronounced reduction in Flipper-TR lifetime (τ) and robust acquisition of SPC⁺ AT2 and RAGE⁺ AT1 fates. In contrast, Matrigel embedding maintained significantly higher CMT and strongly biased organoids toward DP identity (Figure 5A–C).

**Figure 5:**
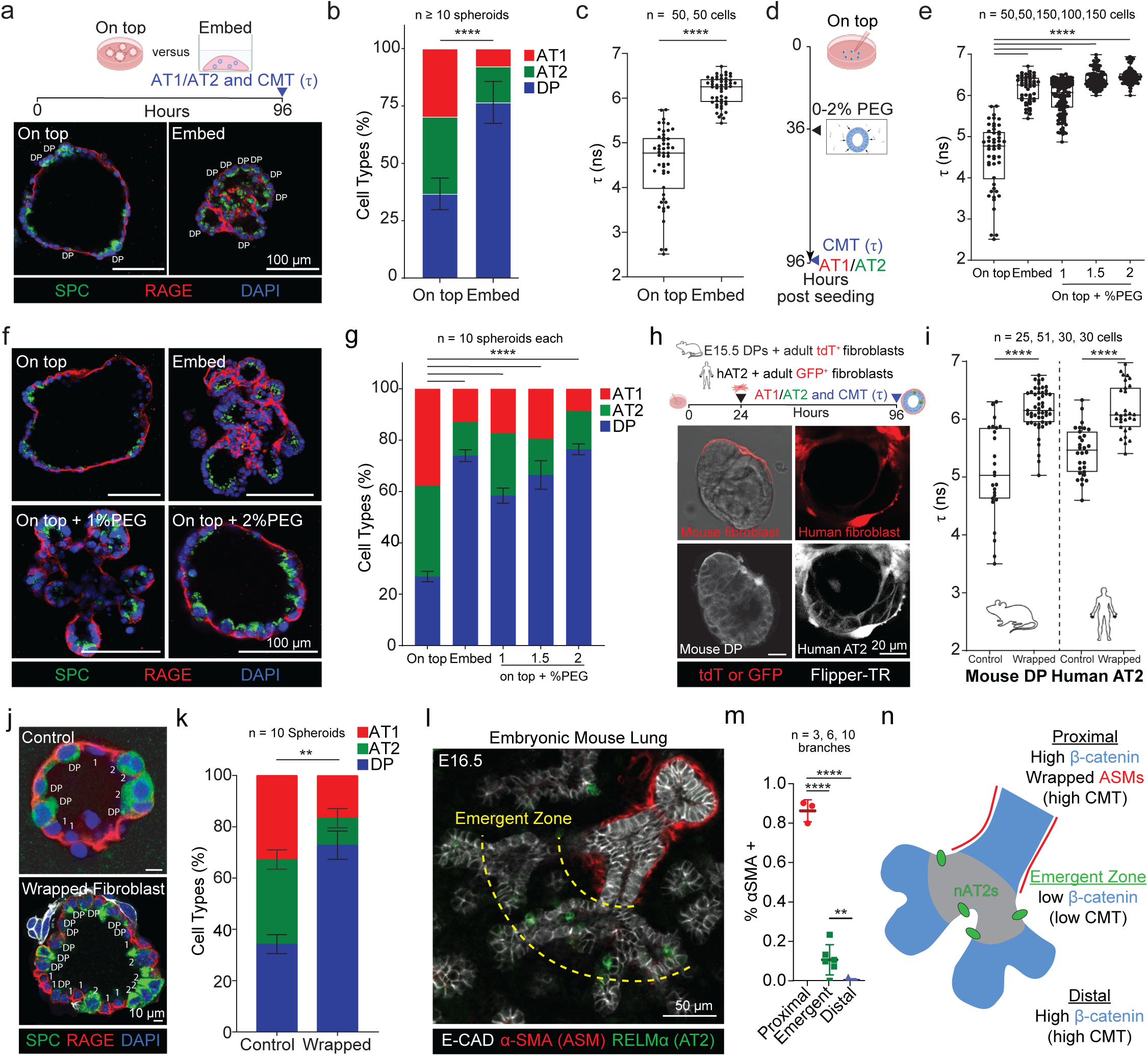
CMT is also regulated extrinsically via direct contact with mesenchymal cells. (A) (Top) Experiment design to determine whether 2D on top versus 3D embed culture impacts CMT and cellular differentiation. (Bottom) Representative alveolospheres from on-top and embed culture stained for SPC, RAGE and DAPI (scale bars, 100µm). (B) Quantification of cell type proportion from conditions in a. Spheroids sampled per condition in experimental duplicate (p-value ****p < 0.0001, Student’s t-test). (C) Quantification of CMT from experiment in A. Data represented as boxplot from 3 independent experiments (p-value ****p < 0.0001, two-tailed Mann-Whitney test). (D) Experiment plan for mimicking isotropic organoid compression to raise CMT via low dose PEG300 in on-top alveolosphere differentiation assay. (E) CMT quantification from both A and D. Data represents mean ±SD from 3 independent experiments (p-values ****p < 0.0001, two-way ANOVA). (F) Alveolospheres cultured as shown in D stained for SPC and RAGE (scale bar, 100µm). (G) Quantification of cell types from F. Data represented as means ± SDs from 3 independent experiments (p-values ****p < 0.0001 calculated by two-way ANOVA). (H) (Top) Experiment design for co-culturing mouse DPs of human AT2s with tdT^+^ adult mouse lung fibroblasts. (Bottom) Representative fibroblast-epithelial spheroids generated at 48 hours wherein epithelial cells can be delineated as present in Flipper-TR and Brightfield but tdT^-^. (I) Quantification of CMT of epithelial cells from H. Data represented as boxplots from 3 independent experiments (p-values ****p < 0.0001, Student’s t-test). (J) E15.5 DPs cultured for 4 days in differentiation media in either Control or fibroblast wrapped condition stained for SPC and RAGE (scale bar, 10µm). (K) Quantification of cell types in J. Data represented as mean ±SD (p-values **p < 0.01, ns p ≥ 0.05, two-way ANOVA). (L) (Left) E16.5 mouse lung stained for α-SMA (airway smooth muscle cells or ASMs), RELMα (nAT2s) and E-cadherin, dashed yellow lines represent the observed emergent zone for nAT2s (scale bar, 50µm). (M) Quantification of α-SMA association in proximal, emergent, and distal regions. (p-values ****p < 0.0001, **p <0.01, Student’s t-test). (N) Schematic of the mode of CMT regulation observed at the proximal (extrinsic and intrinsic), distal (intrinsic), and emergent zone (neither).

To orthogonally elevate extrinsic confinement in a geometry-matched manner, we applied low-dose PEG to on-top cultures to mimic isotropic compression/macromolecular crowding. PEG increased epithelial CMT in a dose-dependent fashion and prevented the CMT reduction observed in control on-top cultures (Figure 5D–E). Consistent with a mechanical constraint on fate acquisition, both Matrigel embedding and PEG treatment suppressed differentiation into AT1 and AT2 cells relative to on-top controls (Figure 5F–G). Together, these results indicate that extrinsic confinement/crowding can maintain elevated CMT and restrict alveolar epithelial differentiation.

We next asked whether analogous extrinsic constraints could arise from epithelial–mesenchymal interactions. Because direct cell–cell contact can elevate CMT in other contexts(Da Silva André and Labouesse, 2024; Li et al., 2021), we established a co-culture assay in which epithelial spheroids (mouse DPs or human AT2s) are wrapped by fluorescently labeled adult lung fibroblasts. Fibroblast co-culture increased epithelial CMT in both mouse and human systems (Figure 5H–I). Moreover, within co-culture spheroids, fibroblast-wrapped configurations exhibited higher CMT than unwrapped controls (Supplemental Figure 4B–C). Functionally, fibroblast wrapping maintained DP-like states and reduced differentiation toward AT1/AT2 fates (Figure 5J–K).

Finally, to relate these findings to in vivo spatial patterning, we quantified αSMA⁺ (airway smooth muscle) association across the E16.5 branching epithelium. Proximal regions displayed extensive αSMA coverage, whereas distal tips and the emergent zone exhibited minimal αSMA association (Figure 5L). Combined with our intrinsic β-catenin–mediated regulation of CMT, these data support a unified mechano-molecular model in which high CMT can be sustained by intrinsic (β-catenin) and/or extrinsic (mesenchymal contact/confinement) inputs, while the emergent zone experiences either constraint the least, permitting CMT reduction and alveolar epithelial differentiation (Figure 5M).

### Two-hit CMT elevation locks adult AT2s in a KRT8⁺ transitional state

Building on our findings that both intrinsic programs (β-catenin–dependent) and extrinsic physical cues (confinement/crowding and mesenchymal contact) can elevate CMT and restrict epithelial fate transitions, we asked whether altered membrane mechanics are a feature of human lung disease. Consistent with this idea, we found that IPF lung tissues exhibit significantly higher epithelial membrane tension compared with non-fibrotic controls (Figure 6A-B). We therefore asked whether sustained exposure to elevated CMT, driven by intrinsic signaling and extrinsic physical constraints, could promote the KRT8⁺ transitional state that accumulates during maladaptive alveolar repair and fibrosis(Choi et al., 2020; Kobayashi et al., 2020; Strunz et al., 2020).

**Figure 6:**
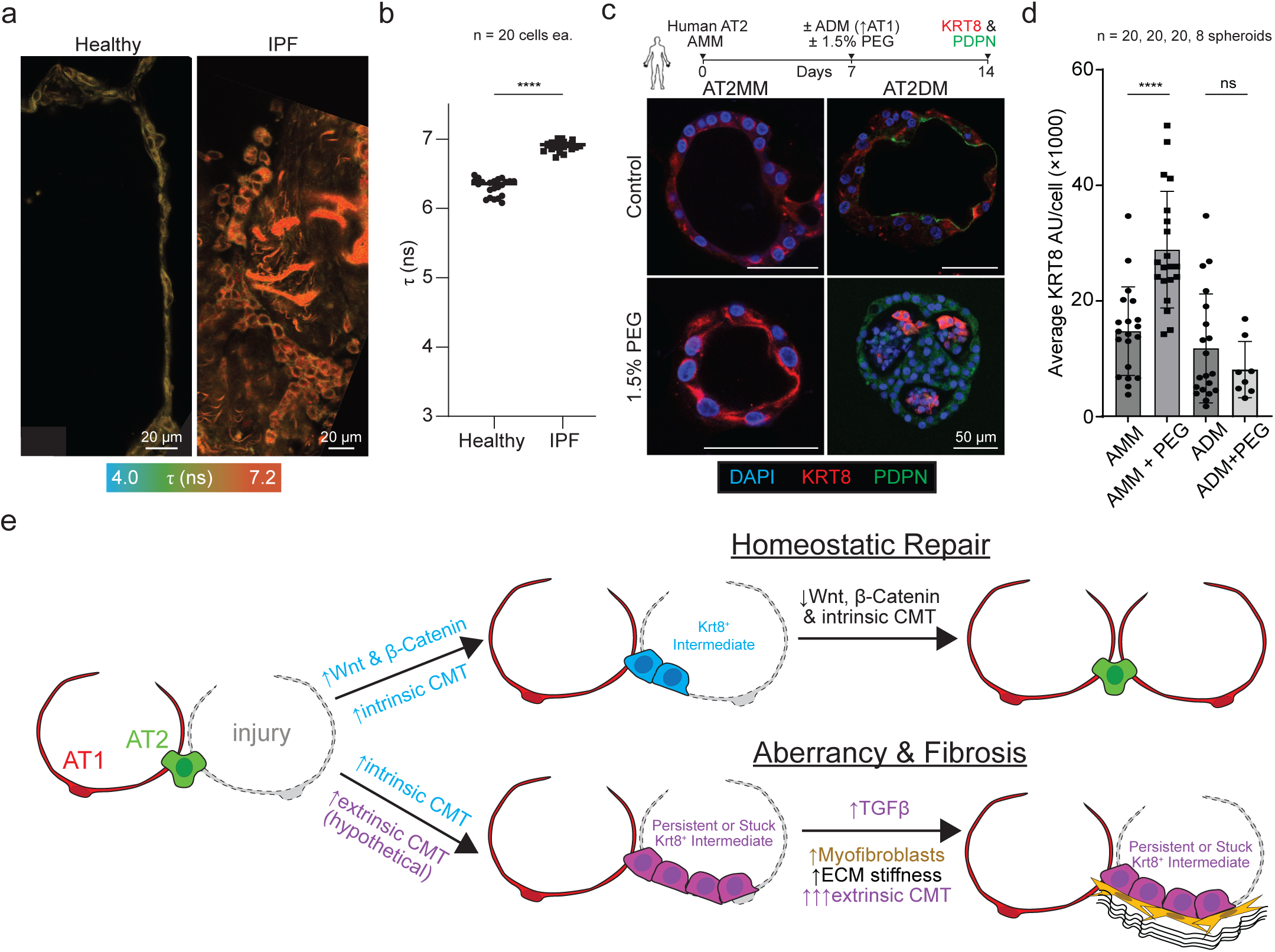
Intrinsic and extrinsic regulation of CMT drives the “stuck” state of AT2 cells during lung injury. (A) Representative Flipper-TR lifetime images of alveolar epithelium in healthy donor and IPF lung tissue. Scale bars, 20 µm. (B) Quantification of fluorescent lifetime of Flipper-TR from control and IPF tissues. (p-values ****p < 0.0001, Student’s t-test). (C) (Upper) Human AT2 organoid cultures. (Lower), Representative images showing KRT8 staining under analogous experimental conditions. (D) Quantification of KRT8^+^ cells (n = 3 independent human donors; mean ± s.e.m.; ****p < 0.0001). (E) Schematic model of intrinsic and extrinsic regulation of CMT in alveolar repair. Intrinsic signals (β-catenin-mediated) and extrinsic mechanical cues (e.g., fibroblast interactions, molecular crowding) converge to modulate CMT. During homeostatic repair, CMT decreases, enabling AT2 differentiation into AT1 cells. In contrast, persistent CMT elevation locks AT2 cells in a “stuck” intermediate state, preventing differentiation and contributing to aberrant repair and fibrosis.

To test this in a controlled setting, we used adult mouse and human AT2 organoids cultured in AT2 maintenance medium (AMM; containing CHIR99021 to stabilize β-catenin) and, in parallel, in AT2 differentiation medium (ADM) to promote AT2-to-AT1 differentiation. In each condition, we imposed an extrinsic crowding/confinement stimulus by adding 1.5% PEG. After 14 days, we quantified KRT8 abundance by immunostaining. In mouse AT2 organoids, PEG significantly increased KRT8 intensity in AMM, whereas PEG did not increase KRT8 in ADM (Supplemental Figure 5A-B). Human AT2 organoids exhibited the same pattern across donors, with robust KRT8 upregulation specifically under AMM+PEG conditions (Figure 6C–D).

Together, these data indicate that extrinsic crowding is not sufficient on its own to induce high KRT8 in differentiating conditions, but instead promotes accumulation of a KRT8⁺ transitional state when combined with a sustained intrinsic β-catenin–stabilized context. We propose a model in which intrinsic (β-catenin–associated) and extrinsic mechanical cues converge to regulate CMT during alveolar repair: under homeostatic conditions, CMT reduction permits AT2-to-AT1 differentiation, whereas persistent elevation of CMT-promoting inputs favors persistence of a KRT8^+^ transitional state and impaired maturation (Figure 6E).

## Discussion

Our findings identify CMT as a central mechanical regulator of alveolar epithelial fate, integrating intrinsic and extrinsic cues to coordinate differentiation, maturation, and repair-associated fate transitions. Across mouse and human developmental tissue and organoid models, we show that CMT decreases prior to—and during—the acquisition of AT1 and AT2 identities, and that preventing this decrease is sufficient to maintain distal epithelial cells in progenitor-like states. Mechanistically, CMT functions as a checkpoint that gates both signaling and architecture: reduced membrane tension permits endocytic activity and enables an FGFR2–ERK program required for AT2 specification(De Belly et al., 2021), while also permitting cytoskeletal–nuclear remodeling that supports YAP nuclear accumulation during AT1 maturation(Riggi et al., 2019; Wu et al., 2017). Conversely, sustained elevation of CMT through combined intrinsic and extrinsic inputs stabilizes undifferentiated or transitional epithelial states and recapitulates key features of the differentiation arrest associated with aberrant repair and fibrosis(Diz-Munoz et al., 2013; Riggi et al., 2019; Wu et al., 2017).

Importantly, CMT reflects membrane–cortex coupling at the cell surface and is distinct from other commonly discussed mechanical parameters such as ECM stiffness or bulk tissue strain(Gokey et al., 2021; Nantie et al., 2018). More broadly, our results highlight membrane tension as a biophysical control point that can tune both trafficking-based signaling and nuclear access, thereby coordinating fate transitions across differentiation systems(Elosegui-Artola et al., 2017). Collectively, these findings position CMT as a mechanical rheostat that couples cell-surface physics to growth factor signaling, cytoskeletal organization, and nuclear architecture to coordinate alveolar fate transitions(Pontes et al., 2017).

During lung development, DPs exhibit high CMT that decreases prior to overt differentiation into AT1 and AT2 fates, and experimentally maintaining high CMT is sufficient to block differentiation in both mouse and human epithelial cultures. These findings align with work in other stem cell systems showing that reductions in membrane tension can be permissive for differentiation, whereas sustained tension stabilizes undifferentiated states(Bergert et al., 2021; De Belly et al., 2021). Our results extend this principle to the alveolar lineage and provide a framework for how membrane mechanics can tune fate decisions in a developing organ.

A key mechanistic link between membrane tension and AT2 fate selection lies in endocytic control of growth factor signaling. FGF signaling is a central determinant of AT2 identity, and receptor trafficking is tightly coupled to downstream signal propagation(Lim et al., 2023; Sun et al., 2022). We find that endocytic activity increases as CMT decreases during late embryonic development, and that experimentally elevating CMT suppresses early endosome abundance and blunts ERK activation. Similar relationships between membrane tension, endocytosis, and signaling have been described in other contexts, where increased membrane tension restricts vesicle budding and receptor turnover(De Belly et al., 2021; Riggi et al., 2019; Wu et al., 2017). Together, our findings support a model in which a developmental reduction in membrane tension relieves an endocytic constraint to enable robust FGF–ERK signaling output and thereby couple the physical state of the cell surface to transcriptional programs driving AT2 fate selection.

CMT also gates a second, distinct module required for alveolar differentiation: cytoskeletal–nuclear remodeling and YAP nuclear accumulation during AT1 maturation. YAP/TAZ signaling is a key determinant of AT1 fate in vivo(Nabhan et al., 2018), and YAP nuclear entry can be regulated by force-dependent transport across nuclear pores(Goodwin et al., 2019; Young et al., 2020a). Recent work has further highlighted the importance of mechanotransductive architecture in AT1 fate, including focal adhesions, actin remodeling, and nuclear–cytoskeletal coupling via the LINC complex(Li et al., 2018; Shiraishi et al., 2023). In our system, emerging AT1 cells exhibit increased nuclear aspect ratio and enhanced nuclear–actin association, and elevating CMT prevents these architectural transitions. Importantly, increased CMT suppresses YAP nuclear accumulation even under enforced YAP expression, consistent with a model in which high membrane tension constrains the architectural remodeling required for effective YAP nuclear access and transcriptional output(Chan et al., 2019). More broadly, these results suggest that membrane mechanics can act upstream of nuclear mechanotransduction pathways by regulating whether cells can adopt the structural states that enable YAP-dependent fate programs(Strunz et al., 2020; Wang et al., 2024).

Our findings also provide insight into how spatial and temporal patterning of alveolar differentiation may be controlled in vivo. We previously described an “emergent zone” between actively proliferating distal tips and airway smooth muscle (ASM)-associated proximal regions where nascent AT2 cells first appear(Sawhney et al., 2025). Here we identify intrinsic and extrinsic mechanisms that can maintain elevated CMT and thereby restrict fate transitions. On the distal side of the branching structure, DPs are enriched for WNT/β-catenin activity, which is often assumed to maintain progenitor status primarily through transcriptional programs(Lim et al., 2023; Nabhan et al., 2018; Sun et al., 2022). Our data suggest an additional, underappreciated mechanism: β-catenin can stabilize a high-CMT state through a transcription-independent function consistent with its junctional role, likely by reinforcing membrane–cortex attachment via adherens junction organization(Borghi et al., 2012; Desai et al., 2009; Valenta et al., 2012). In support of this model, stabilizing β-catenin elevates CMT and blocks fate transitions even when β-catenin–dependent transcriptional output is inhibited, while genetic depletion of β-catenin reduces CMT and promotes AT1-like transitions in adult AT2 cultures—consistent with prior work linking WNT/β-catenin perturbations to AT2 fate maintenance(Nabhan et al., 2018).

On the proximal side, epithelial branches are physically associated with ASM. Although the exact role of ASM remains unclear in epithelial branching morphogenesis(Goodwin et al., 2019; Young et al., 2020b), its intimate contact with proximal epithelium raises the possibility that mesenchymal wrapping provides an extrinsic, spatially patterned constraint on epithelial state transitions. Consistent with this idea, we developed an epithelial–mesenchymal co-culture model and found that direct mesenchymal contact elevates epithelial CMT and biases epithelial cells toward progenitor-like states in both mouse and human organoid systems. Direct cell–cell adhesion has also been shown to elevate membrane tension in other contexts, supporting the plausibility of a contact-mediated mechanical constraint(Bijonowski et al., 2025; Borghi et al., 2012). Together, these findings support a unified “mechano-molecular” model in which alveolar differentiation is favored in regions where neither intrinsic (β-catenin-associated) nor extrinsic (mesenchymal contact/confinement) inputs maintain high CMT, thereby creating a permissive window for fate acquisition(Sawhney et al., 2025; Zepp et al., 2017).

Beyond in vivo patterning, our findings emphasize that CMT is sensitive to biophysical cues from the cellular microenvironment. Matrix embedding and isotropic compression/macromolecular crowding elevated CMT and inhibited differentiation in vitro, indicating that confinement alone can be sufficient to maintain a high-tension, progenitor-like state(Chan et al., 2019; Li et al., 2021). This has practical implications for organoid-based differentiation systems: culture geometry and crowding can impose mechanical constraints that alter endocytosis-dependent signaling(Sorkin and von Zastrow, 2009) and fate outcomes(Konishi et al., 2022; Li et al., 2021), and should be considered when interpreting organotypic models of development or repair.

Finally, our work suggests a testable link between membrane mechanics and maladaptive epithelial repair in fibrosis. A KRT8^+^ transitional state has been observed during alveolar regeneration and is reported to persist in human pulmonary fibrosis(Choi et al., 2020; Jiang et al., 2020; Kobayashi et al., 2020; Strunz et al., 2020). In adult mouse and human AT2 organoids, sustained exposure to an intrinsic β-catenin–stabilized context together with extrinsic crowding promotes a robust increase in KRT8 abundance, consistent with accumulation of a transitional state under conditions that, in our system, elevate CMT(Choi et al., 2020; Kobayashi et al., 2020; Strunz et al., 2020). These observations raise the possibility that pathological maintenance of high membrane tension—driven by combined intrinsic programs and extrinsic microenvironmental constraints—represents a barrier to terminal AT2-to-AT1 differentiation following injury(Kobayashi et al., 2020; Strunz et al., 2020; Wang et al., 2023; Wang et al., 2024). Future studies will be important to determine whether CMT is elevated endogenously during fibrotic remodeling, which extrinsic cues most strongly impose high CMT on alveolar epithelium, and whether lowering CMT can restore effective re-epithelialization(Wang et al., 2023; Wang et al., 2024).

Collectively, our findings establish CMT as a unifying mechanical signal that integrates molecular, cellular, and tissue-scale inputs to govern alveolar epithelial fate. By acting as a physical checkpoint that gates both receptor trafficking and nuclear mechanotransduction, CMT provides a mechanism for coupling the biophysical state of the cell surface to lineage-specific transcriptional programs(Diz-Munoz et al., 2013; Elosegui-Artola et al., 2017; Pontes et al., 2017; Sorkin and von Zastrow, 2009). We propose that developmental modulation of CMT defines a window of competence for alveolar differentiation, while persistent elevation of CMT contributes to repair failure by stabilizing transitional epithelial states associated with fibrosis(Choi et al., 2020; Jiang et al., 2020; Kobayashi et al., 2020; Strunz et al., 2020).

## Study Limitations

We briefly note limitations that motivate future work. First, much of our mechanistic dissection relies on organoid, matrix-context, and epithelial–mesenchymal co-culture systems, which capture key aspects of alveolar fate transitions but cannot fully reproduce in vivo geometry, ventilation-driven stretch, vascular interfaces, or inflammatory cues. Second, our CMT readouts (FLIM-based membrane probe lifetime and optical-tether forces) report effective membrane mechanics but do not fully resolve contributions from lipid bilayer tension versus membrane–cortex attachment, nor do they capture rapid or subcellular fluctuations. Third, pharmacologic approaches used to modulate CMT or downstream pathways (e.g., cholesterol extraction/crowding and dynamin/ERK/YAP pathway perturbations) may have off-target effects; complementary genetic and orthogonal biophysical manipulations will be important. Finally, the “stuck” KRT8⁺ transitional state is modeled by combined intrinsic and extrinsic CMT elevation in culture and thus may not recapitulate the full cellular heterogeneity or immune milieu of fibrotic lungs, underscoring the need to test whether CMT is elevated in vivo during injury/fibrosis and whether lowering CMT improves repair.

## Methods

### Ethics

All mouse experiments followed applicable regulations and guidelines and were approved by the Institutional Animal Care and Use Committee at the Mayo Clinic (Protocol# A00006319-21). Mice were housed under a 12-hour light/12-hour dark cycle and lights were not used during the dark cycle. Mouse housing temperature was kept at 65-75°F (∼18-23°C) and humidity at 40–60%. Mice were housed in filtered cages, and all experiments were performed in accordance with approved Institutional Animal Care and Use Committee protocols and ethical considerations at the Mayo Clinic. Human AT2 cells, lung fibroblasts and slices were obtained from a Mayo Tissue Biobank repository (IRB #24-012991). Human lung tip progenitors were isolated from tissues collected under IRB #25-004599.

### Mouse strains

Timed-pregnant C57BL/6 J females (abbreviated B6; Jackson Laboratories) were used for all embryonic time points, with gestational age verified by crown–rump length. For studies of adult wild-type lungs, B6 males and females were used. Mosaic labeling and deletion studies were conducted by Cre recombinase expression using gene-targeted alleles Nkx2.1-Cre; mTmG (*Tg^Nkx2-1-cre^; Rosa26^mTmG^),* Sftpc-CreER; mTmG (*Sftpc^cre/ERT^; Rosa26^mTmG^)*, YAP-Ctrl (*Tg^Nkx2-1-cre^*; *Tigre^Ai140^*), YAP-OE (*Tg^Nkx2-1-cre^*; *_Tigre_^Ai140^_; Rosa26_^teto-S112A-YAP1^_)_*_, β-catenin_^fl/fl^ _(*Ctnnb1*_*^flox/flox^*). Genomic DNA was extracted from ear punches followed by Proteinase K (KAPA Biosystems) digestion, and genotyping was performed by polymerase chain reaction (PCR) using published primer sets.

### Mouse Cell isolation and culture

Adult AT2 cells and embryonic (E15.5, E16.5 and E17.5) distal progenitors were purified as described previously(Ali et al., 2021; Brownfield et al., 2022). Adult 2-month-old mice and pregnant mice were euthanized by the administration of CO_2_. For E15.5, E16.5 and E17.5 embryos were removed from the mother, and their lungs were isolated en-bloc without perfusion and pooled by litter (5 to 7 embryos) and kept in cold phosphate buffered saline (PBS, pH 7.4). Lungs were microdissected to remove the proximal lung tissue, leaving only the distal tissue. Distal lung tissues were dissociated in 5U/ml dispase (Stemcell technologies). Tissue suspension was gently triturated with different diameter sized Pasteur pipettes until a single cell suspension was reached. For adult lungs, the vasculature was perfused through the right ventricle with chilled PBS. The trachea was punctured, and lungs were inflated with 50U/ml dispase (Corning) for 45 minutes at 25°C. Digested lungs were minced and triturated briefly with a 5ml pipette.

To deplete red blood cells, an equal volume of DMEM/F12 supplemented with 10% FBS and 1U/ml penicillin-streptomycin (Thermo Scientific) was added to the lung single cell suspensions before filtering through 100µm mesh (Fisher) and centrifuging at 400×g for 10 minutes. Pelleted cells were resuspended in red blood cell lysis buffer (BD biosciences), incubated for 2 minutes, passed through a 40µm mesh filter (fisher), centrifuged at 400×g for 10 minutes, and then resuspended in MACS sorting buffer (Miltenyi Biotec).

Bipotent progenitors from E15.5 lungs were isolated and enriched from the single cell suspensions by using MACS Microbeads technology with MS columns (Miltenyi Biotec #130-042-201). Antibodies against CD31, CD45, Pdgfrα, Ter119 (Miltenyi Biotec #130-097-418; #130-052-301; #130-101-502; #130-049-901) were used to magnetically deplete positive cells. For positive selection of cell types, the following antibodies were used: anti-EpCAM (clone G8.8, eBioscience, 13-5791-82) for E15.5 distal bipotent progenitors, anti-RELMα (PeproTech, 500-P214BT) for E16.5 and E17.5 nascent AT2 cells. Following antibody incubation, the cells were then incubated with anti-biotin microbeads (Miltenyi Biotec #130-090-485) and loaded on MS columns as per vendor’s protocol. The flow through was discarded, and the column was taken off the magnet. The remaining cells in the column were eluted with the MACS buffer. Adult AT2 cells were isolated from single-cell suspensions of adult lungs in the same way for positive selection of MHC Class II (clone M5/114.15.2, eBioscience, 13-5321-82) positive AT2 cells, generating a highly enriched preparation of AT2 cells. To enrich adult fibroblasts, MACS depletion was conducted as above using antibodies against CD31, CD45, and EpCAM.

Mouse lung fibroblasts were isolated from *Rosa26^mTmG^*mice lacking Cre recombinase. Lungs were dissociated as described above for adult AT2 isolation, and single-cell suspensions were incubated in MACS isolation buffer (Miltenyi Biotec, Cat#130-091-221) with antibodies against CD45, CD31, and EpCAM for 15 min. Cells were centrifuged at 400 × g for 5 min, resuspended in 1 mL buffer, and passed through an LD column. Flow-through cells were plated and allowed to adhere overnight. Non-adherent cells were removed by washing the following day, leaving tdTomato⁺ fibroblasts. Fibroblasts were cultured in DMEM supplemented with 10% FBS and 1% penicillin–streptomycin, with media changes every 72 hours and passaging at ∼90% confluence.

### Mouse primary cell culture

For culturing bipotent progenitors, cell density and viability were calculated using a Vi-CELL XR cell viability analyzer (BD). A density of 30,000 cells per well was used for culture in 8-well #1.5 coverglass chambers (Cellvis, C8-1.5H-N), pre-coated with growth factor reduced Matrigel (80μl, Corning 354230) for 30 minutes at 37°C. Cells were supplemented with 50ng/ml FGF7 (R&D Systems), 1mM Methyl-β-cyclodextrin (Sigma C4555), 10µM Dynasore (Tocris #2897), 10µM FR180204 (Tocris #3706), 10μM DAPT (15020 Cell signaling), 5µM Akti-1/2 (Tocris #5773), 3nM KRpep-2D (Selleckchem #S8499), 3μM PLC **γ** (Tocris # U73122), 50μM NSC 74859 (Tocris # 4655), 3µM CHIR99021, 10μM TRULI (HY-138489), and 100 nM Triptonide (HY-32736) in DMEM/F12 as required for various experiments. Cells were maintained at 37°C in 400μl of DMEM/F12 with media changes every other day in a 5% CO_2_/air incubator typically for four days, except where indicated otherwise.

For embed culturing, 30,000 DP cells mixed in matrigel and then plated in 8-well #1.5 cover glass chambers (Cellvis, C8-1.5H-N). Cells were maintained at 37°C in 400μl of DMEM/F12 supplemented with 50ng/ml FGF7, with media changes every other day in a 5% CO2/air incubator for four days. For osmotic stress experiments, hyperosmotic stress was applied by adding polyethylene glycol 300 (PEG300) from MedChem Express (HY-Y0873) to isotonic culture medium(Li et al., 2021). Cells cultured on top of Matrigel in DMEM/F12 media supplemented with 50ng/ml FGF7. After 36 hours, the media were changed to DMEM/F12 + 50ng/ml FGF7 supplemented with following concentrations of PEG300; 1%, 1.5%, and 2%. Cells were maintained for another 50 hours in a 5% CO2/air incubator.

For culturing adult AT2 cells, cells were seeded in AT2 expansion media as described previously(Konishi et al., 2022) for 7 to 10 days.

### Human Cell isolation and culture

Primary human AT2 cells were isolated from distal lung tissue obtained from donor lungs not used for transplantation, in accordance with institutional ethical guidelines and with informed consent. AT2 cells were isolated using established epithelial enrichment and sorting strategies as previously described(Konishi et al., 2022). Purified AT2 cells were expanded on-top of Matrigel in 8-well chamber slide wells coated with Matrigel using a published protocol for feeder-free organoid-expansion(Konishi et al., 2022). Media was replaced every 2–3 days, and organoids were passaged as described in the original protocol. AT2 cell identity was confirmed by expression of canonical markers, including SFTPC and HTII-280, prior to downstream experiments.

Primary human lung fibroblasts were isolated from distal lung tissue obtained from the same donor sources. Lung tissue was enzymatically digested and plated under standard adherent culture conditions to allow selective attachment and expansion of fibroblasts in DMEM with 10% FBS and 1% Penicillin/Streptomycin solution. Fibroblasts at 60% confluency were treated with supernatant from GFP^+^ lentivirus-producing HEKs in serum-free media overnight using 100x ABM transduction enhancer (G515) before having their normal culture media restored the next morning.

### Human lung slice culture and Flipper-TR Imaging

Human lung PCLS were derived from explanted pieces of lungs or whole lungs through IRB #24-012991. For healthy tissue, a whole transplant-rejected lung was received. Single lobes were cordoned off and inflated with 50-200 mL of 2% LMP agarose. When the tissue was sufficiently stiffened by agarose, 5 mm biopsy punches were used to make biopsy cores for use on the compresstome, which were halved lengthwise before being used for PCLS creation. For IPF tissue, tissue was received as small pieces (1-3 cm^3^) from either healthy or ILD conditions, including IPF. Samples were injected with 2% LMP agarose through any visible airways; if possible, 5 mm biopsy cores were used to make tissue punches. Once prepared, all samples were then glued into the slicing apparatus of a standard compresstome, surrounded by 2% LMP agarose, chilled with a freezing block, and sliced to 500 µm thickness. Given the inconsistent inflation of tissue pieces, slices were not always of consistent thickness – samples were selected for analysis based on approximately correct thicknesses and general appearance.

For membrane tension measurements, samples were incubated with 1μM Flipper-TR in serum-free culture medium for 10 minutes at 37°C in a humidified atmosphere containing 5% CO₂ prior to imaging. FLIM data were acquired using standard time-correlated single-photon counting (TCSPC) techniques, with excitation provided by a 488 nm pulsed laser and photon detection through a 600 ± 50nm bandpass emission filter.

### Human fetal lung progenitor cell isolation, expansion and differentiation

Human fetal lung tissue was obtained from legally consented specimens in compliance with IRB approval and all applicable institutional and regulatory guidelines. Distal lung epithelial progenitor cells were isolated as previously described and expanded following established methods developed by Emma Rawlins and colleagues(Lim and Rawlins, 2024). Briefly, epithelial progenitor cells were cultured on top of Matrigel and maintained in defined progenitor expansion medium that supports distal lung epithelial identity and self-renewal. Culture medium was replaced every 2–3 days, and cells were passaged according to the published protocol.

For differentiation assays, progenitor cells were maintained under expansion conditions for 14 days, after which the culture medium was switched to alveolar type 2 (AT2) differentiation medium, with or without PEG300 supplementation, for an additional 7 days prior to downstream analyses. Progenitor identity was confirmed by expression of distal epithelial markers, including SOX9, and AT2 differentiation was assessed by induction of SFTPC expression.

### Plasmids

All plasmids used in this study were either obtained from commercial sources or generated using standard molecular biology techniques. pMAX-GFP (GFP control) was provided with Lonza nucleofection kits. MEK1-GFP and ERK1-GFP plasmids were obtained from Addgene (plasmid #14746 and #14747, respectively). The membrane-targeted mCherry construct (Mem-mCherry) was derived from Addgene plasmid #55779 and subcloned into the pMAX-GFP backbone using Gibson Assembly (New England Biolabs, #E2621S). The iMC-linker construct was obtained from a plasmid provided by the Dis-Muñoz laboratory(Diz-Munoz et al., 2013) and subcloned into the same CMV-driven pMAX-GFP backbone. Plasmids were prepared using endotoxin-free maxi-prep kits according to the manufacturer’s instructions. All plasmids were sequence-verified prior to use.

### Nucleofection of primary cells

Distal lung progenitor cells were centrifuged and resuspended at a density of 5 × 10⁶ cells/mL in Lonza nucleofection buffer. Aliquots of 20µL cell suspension were transferred to individual tubes and mixed with experimental or control plasmid DNA (1µg per nucleofection in TE buffer). Cell–DNA mixtures (20µL) were loaded into a 16-well nucleofection cuvette, taking care to avoid bubble formation. Nucleofections were performed using a Lonza 4D-Nucleofector system with P3 Primary Cell Nucleofector Solution (Lonza) and program CM-113.

Immediately following nucleofection, cells were transferred to ice-cold nucleofection recovery buffer and seeded onto Matrigel-coated culture surfaces in recovery medium consisting of alveolar expansion medium lacking CHIR99021. After 36 hours, the medium was replaced with DMEM/F12 supplemented with FGF7 (50ng/mL) and penicillin–streptomycin, and cells were cultured for an additional 36 hours.

### Co-culturing protocol

10,000 DP cells per well were added to a cover-slip bottom 96 well plate treated to allow control over adhesion and grown in DMEM/F12 media supplemented with 50ng/ml FGF7 and 2% matrigel. After 24 hours, DP cells were mixed with tdTomato labeled fibroblasts in 1:1 ratio and maintained in media for 48 hours before taking live Flipper-TR measurements for membrane tension. After membrane tension measurements, cells were fixed in 4% PFA for further immunostaining.

### Mosaic deletion of β-Catenin

To mosaically delete β-Catenin in adult AT2 cells, we used a recombinant AAV described below to infect and constitutively express Cre recombinase to delete the conditional β-catenin^fl/fl^ allele. The pAAV.CMV.HI.eGFP-Cre.WPRE.SV40 was a gift from James M. Wilson (Addgene viral prep # 105545-AAV9; http://n2t.net/addgene:105545; RRID: Addgene_105545) purchased directly from Addgene. β-catenin^fl/fl^ AT2 expanded to form spheroids (size ≥ 100µm) for 7 to 10 days in AT2 expansion media. Before adding viral particles, cells are treated with ABM transduction enhancer (G515) for 10 minutes, followed by 25µL of 1×10^12^ vg/mL virus added to the culture media. Cells were maintained in expansion media for another 4 days for taking Flipper-TR measurements followed by fixation and staining. Cells positive for pAAV.CMV.HI.eGFP-Cre.WPRE.SV40 showed nuclear GFP expression and were selected for Tau gating measurements.

### Lentivirus and Adeno-associated Virus Creation

Lenti-X HEK cells were plated on 10cm cell-culture treated dishes. At 70% confluency, cells were transfected with 2nd-generation lentiviral plasmids generated by the Trono Lab (PsPax2, 12260, Vsv-g, 8454) and FUGW-GFP (14883). For transfection, plasmids were mixed in 1mL serum-free DMEM in a 1:1:1 molar ratio and totaling 10µg. 30µL of 1mg/mL PEI solution was added, and the solution was vigorously vortexed for 15 seconds before being allowed to sit for 20 minutes at room temperature. The plasmid solution was added to the HEK cells, and the media changed to fresh serum-free DMEM the next day. Lentiviral supernatant was collected at 48- and 72-hours post-transfection and used to treat human and mouse fibroblasts.

Adeno-associated virus (AAV) vectors were produced in HEK293T cells using a similar system, followed by purification using a chloroform extraction method of cell lysate as previously described. Briefly, cells were co-transfected with plasmids encoding the AAV vector genome, packaging components, and helper plasmid. Virus-containing cells and media were harvested, lysed, and subjected to chloroform-based extraction to remove cellular debris and contaminants. The aqueous phase containing AAV particles was clarified and concentrated before addition to distal progenitor cells.

### Lung isolation and processing

For prenatal time points, individual embryos were staged by fetal crown–rump length prior to sacrifice, and lungs were removed en bloc. For all time points, lungs were separated by lobe and fixed in 4% PFA at 4 °C for 1 (E15.5–E16.5), 1.5 (E17.5), or 2 (E18.5–P10) hours. Fixed tissues were washed in PBS and dehydrated in increasing methanol graded and stored in 100% methanol at -20°C until further processing(Gillich et al., 2021). For sectioning, lungs were rehydrated and embedded as required and sectioned using a vibrating microtome (Leica VT1000S) to generate tissue sections of uniform thickness (50–100µm).

For live imaging experiments, E15.5, E16.5, and E17.5 lungs were removed en bloc, washed in ice-cold PBS, and embedded in agarose at 4 °C for 30 min prior to sectioning. Live tissue sections were collected in ice-cold RPMI-1640 medium and mounted in 2-well #1.5 coverglass chambers for in situ Flipper-TR–based membrane tension measurements.

### Immunostaining

Immunohistochemistry was performed as previously(Brownfield et al., 2022) described with minor modifications. Fixed spheroids and lung sections were permeabilized and blocked in blocking buffer consisting of 5% goat serum in PBS containing 0.5% Triton X-100 at 4 °C. Primary antibodies were used at a dilution of 1:200 for PFA-fixed spheroids and 1:500 for tissue sections unless otherwise noted. The following primary antibodies were used: rabbit anti pro–Sftpc (Sigma-Aldrich), rabbit anti SPC (Invitrogen), rat anti RAGE (R&D Systems), rat anti E-cadherin (Life Technologies, ECCD-2), hamster anti podoplanin (DSHB, 8.1.1), hamster anti Mucin1 (Thermo Fisher Scientific, HM1630), mouse anti FGFR2 (Santa Cruz Biotechnology), rabbit anti EEA1 (Cell Signaling Technology), rabbit anti β-catenin (Cell Signaling Technology), anti-mouse phalloidin-647 conjugated (Abcam), mouse anti YAP (Santa Cruz Biotechnology), mouse anti α-smooth muscle actin IgG2a (Invitrogen), rabbit anti RELMα (PeproTech), mouse anti SOX9 (Invitrogen), rabbit anti SOX9 (Millipore), rat anti SOX2 (Invitrogen), Sheep anti podoplanin (R&D Systems), and mouse anti TGFBR2 (Proteintech).

Following washes, samples were incubated with Alexa Fluor–conjugated secondary antibodies (1:1000 dilution unless otherwise noted), including goat anti-rabbit Alexa Fluor 555 (Invitrogen, A21428), goat anti-rabbit Alexa Fluor 647 (Invitrogen, A32733), goat anti-rat Alexa Fluor 488 (Invitrogen, A11006), goat anti-rat Alexa Fluor 647 (Invitrogen, A21247), goat anti-hamster Alexa Fluor 647 (Invitrogen, A21451), and goat anti-hamster Alexa Fluor 594 (Invitrogen, A21113). Samples were mounted in VECTASHIELD Antifade Mounting Medium (Vector Laboratories, H-1000) and sealed with coverslips. Samples were stored at 4°C in the dark.

### Fluorescence microscopy and FLIM analysis

For both in vitro and in situ measurements, alveolospheres and live precision-cut lung slices (PCLS) were maintained on glass-bottom multi-well plates. Fluorescence lifetime imaging microscopy (FLIM) of cultured alveolospheres and live PCLS was performed using an inverted Leica Stellaris 8 microscope equipped with environmental control to maintain 37°C and 5% CO₂.

For membrane tension measurements, samples were incubated with 1μM Flipper-TR in serum-free culture medium for 10 minutes at 37°C in a humidified atmosphere containing 5% CO₂ prior to imaging. FLIM data were acquired using standard time-correlated single-photon counting (TCSPC) techniques, with excitation provided by a 488 nm pulsed laser and photon detection through a 600 ± 50nm bandpass emission filter.

Photon arrival histograms were extracted from regions of interest (ROIs) or individual pixels and fitted using a double-exponential decay model to obtain two lifetime components (τ₁ and τ₂). For FLIM analyses shown in Figures. 1, 3, 4, and 5, data were fitted using the Leica LAS X software according to a two-exponential reconvolution model. The longer lifetime component (τ₁), corresponding to the decay component with the higher amplitude, was used to report membrane tension. τ₁ values ranged from approximately 2.0 to 7.2ns, with longer lifetimes indicating higher membrane tension, as previously described(Colom et al., 2018).

### Confocal imaging and quantitative image analysis

Confocal images were acquired using the Leica Stellaris 8 microscope and analyzed using ImageJ software (National Institutes of Health). Mean fluorescence intensity (MFI) measurements for EEA1, FGFR2, YAP, and β-catenin staining in histological preparations were quantified using the mean fluorescence method in ImageJ and normalized to the area of the corresponding region of interest (ROI).

Alveolosphere cell-type composition was quantified by manual cell counting from confocal images of histological sections using ImageJ. Cells were classified as AT2 cells when positive for pro–Sftpc, AT1 cells when positive for RAGE, and DP cells when positive for both pro-Sftpc and RAGE. Quantification was performed on a per-cell basis across experimental conditions.

### Tether pulling and trap force measurements

Trap force measurements were performed using a home-built optical tweezer system incorporating a 4 W, 1064nm laser (Ventus, Laser Quantum) coupled to an inverted microscope (Nikon Eclipse TE2000-U) equipped with a 100× oil-immersion objective (NA 1.30, CFI Plan Fluor DLL, Nikon) and a motorized stage (PRIOR ProScan). Optical trap calibration was performed as previously described(Han et al., 2018).

Carboxylated latex beads (1.9µm diameter; Thermo Fisher Scientific, C37278) were coated with concanavalin A (50µg/mL) by incubation on a shaker for 1 h prior to experiments. Bead position was recorded in brightfield at 90ms intervals before and during membrane tether formation. Trap forces were calculated from bead displacement using the calibrated trap stiffness and bead position, analyzed with a custom Fiji plugin (HDB; Schindelin et al., 2012)(Schindelin et al., 2012). Typical trap stiffness values were ∼0.13pN/nm.

### Fluid phase uptake assay

Cells were seeded on IBIDI dishes (IBIDI Scientific, 81156) a few hours before imaging. Cells were then incubated for 10 minutes in DMEM/F12 media with 10mg of pH-Rhodo Dextran (pHrodo Red Dextran, 10,000 MW, for Endocytosis, Thermo Fisher Scientific). Cells were then rinsed immediately prior to imaging. Each dish was imaged for 30 minutes. For quantifications of pH Rhodo, total intensity inside the cell was measured with Fiji by using a sum z projection. The signal was corrected by taking a ROI of the background of a similar size as of the measured cells. The background signal was then subtracted from the signal from cell. Only single cells or groups of cells with clear boundaries were quantified. Experiments were always performed side by side with cells in DMEM/F12 media as control.

### Statistical analysis

Data analysis and statistical tests were performed with R or GraphPad Prism software. Replicate experiments were all biological replicates with different animals, and quantitative values were presented as mean ± SD unless indicated otherwise. Two-sided Student’s t-tests (for normally distributed data), Mann-Whitney U-tests (for data not normally distributed), and One-way ANOVA (for comparisons between ≥ 3 groups) were used to determine P-values. No statistical method was used to predetermine sample size, and data distribution was tested for normality prior to statistical analysis and plotting. Statistical significance was defined as p < 0.05 unless otherwise stated. Both male and female animals were included in experiments, and biological replicates and group comparisons were age- and sex-matched where applicable.

### PROGENy analysis

For analysis of lung development through embryonic, perinatal, and adult timepoints in published scRNAseq datasets, the processed mRNA counts for each cell were used from developing mouse lung dataset GSE149563. It was processed into identifiable clusters and annotated as described previously.(Sawhney et al., 2025) Single-cell signaling pathway activities were inferred using the PROGENy R (1.28.0) package. The top 100 responsive genes per pathway were used to calculate activity scores from the log-normalized expression matrix, added to the metadata of the Seurat object, and plotted using the DimPlot function.

---

**Supplementary Figure 1:**
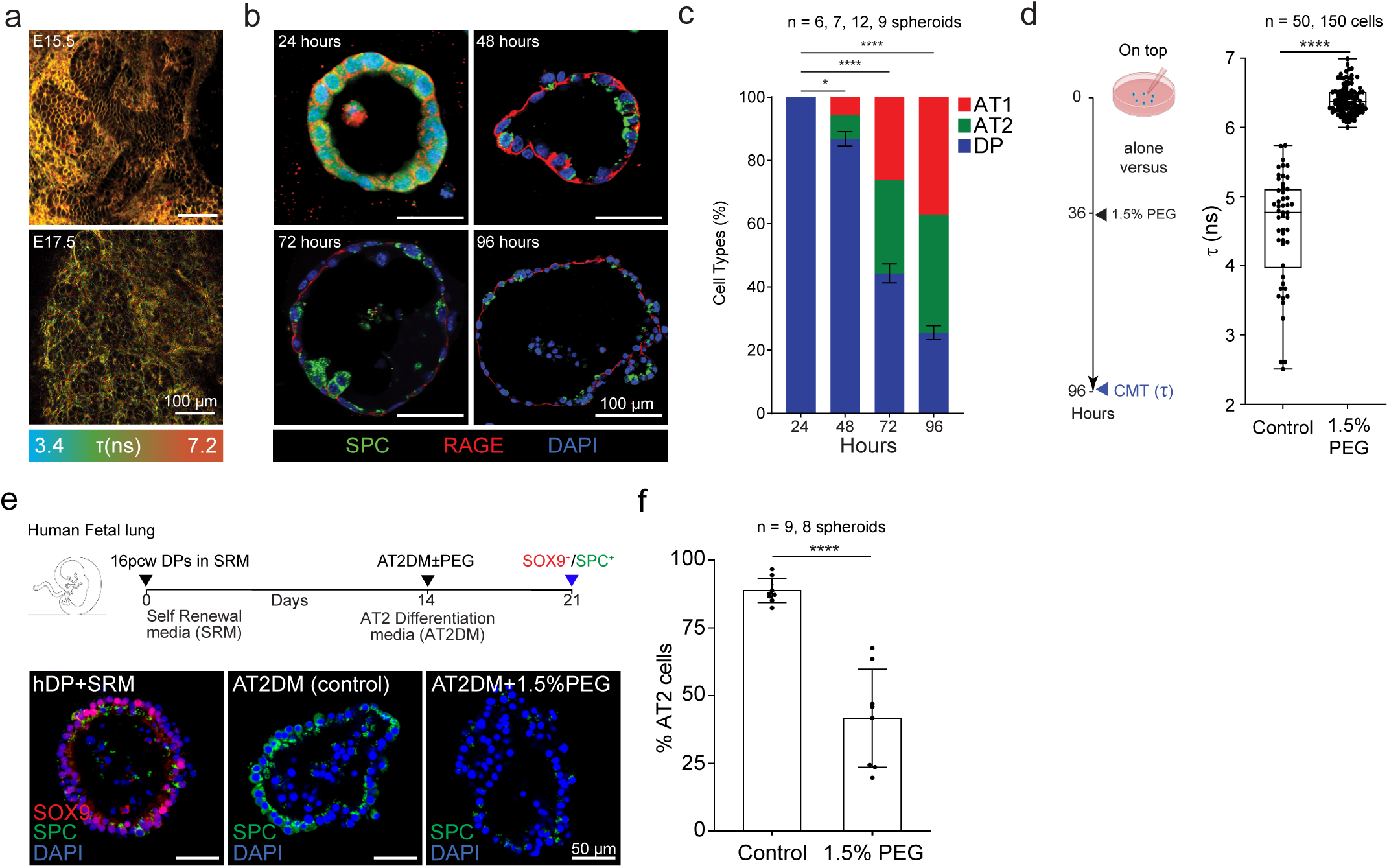
(A) Representative images of embryonic lung tissues at stages E15.5 and E17.5 stained with Flipper-TR. The scale bar represents 100µm. (B) Immunostained images show DP cultured on Matrigel for four days, with media supplemented every two days with 50ng/ml FGF7. Fixation occurred at 24-hour intervals: 24, 48, 72, and 96 hours. The fixed spheroids were stained for AT1 marker Rage (red), AT2 marker Sftpc (green), and nuclear DAPI (blue). Scale bars indicate 100µm. (C) Graph showing the percentage of DP cells per spheroid at 24-hour intervals (n = 10 spheroids for each condition, in triplicate). p values (*0.0145, **** <0.0001) were calculated using one-way ANOVA with multiple comparisons. (D) DP cell seeded on matrigel supplemented with 50ng/ml FGF7. Media was supplemented with 1.5% PEG300 to raise CMT. CMT was measured by τ-gating of PEG-treated spheroids vs control after 4 days. Data are shown as means ± SDs from 3 independent experiments. p value (****<0.0001) was calculated using a Mann-Whitney test. n = 50 and 100 cells for control and 1.5% PEG groups, respectively. (E) Above: Schematic demonstrating protocol used for expansion and differentiation of human fetal lung distal lung tip progenitor cells. Below: Representative images of expanded cells, cells treated with AT2 differentiation media (AT2DM), and AT2DM + 1.5% PEG. (F) Quantification of % of Sftpc+ cells in AT2DM(Control) and AT2DM + 1.5% PEG conditions. P-value (**** < 0.0001) was calculated using a Mann-Whitney test.

**Supplemental Figure 2:**
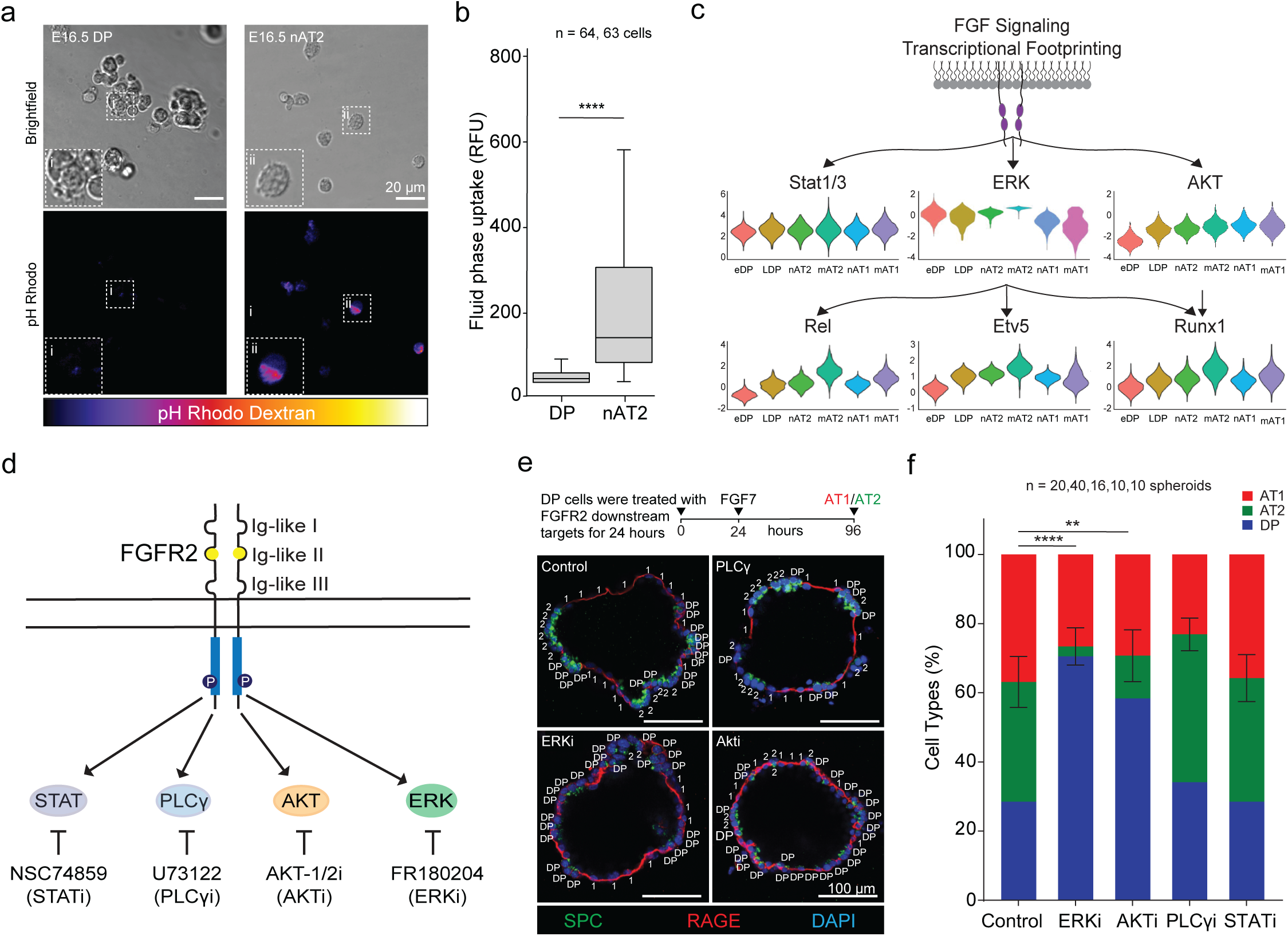
(A) Representative images of pH Rhodo Dextran dye uptake assay of E16.5 DPs and nAT2 cells. (B) Quantification of fluid phase uptake by DP and nAT2. Statistical significance was determined using Student’s t-test yielding p-values of < 0.0001. (C) Transcriptional footprinting analysis of FGF signaling by using scRNA seq data of alveolar epithelium. (D) Schematic diagram of Fgfr2 downstream signaling target and their inhibition with small molecules. (E) Upper, timeline for mouse DPs treated with small molecule inhibitors for 24 hours and then culture in FGF7 for another 72 hours before fixation and staining for AT1 and AT2 markers at 96 hours. Lower, representative images of spheroids treated with small molecule inhibitors vs control. (F) Quantification of spheroids in (E) demonstrate a significant reduction in AT2 differentiation after ERK inhibition. Statistical significance was determined using one-way ANOVA yielding a p-value of ****< 0.0001 and **<0.001. n = 20,40,20,20 and 20 for the Ctrl, Erki, AKTi, PLCγi, and STATi groups respectively.

**Supplemental Figure 3:**
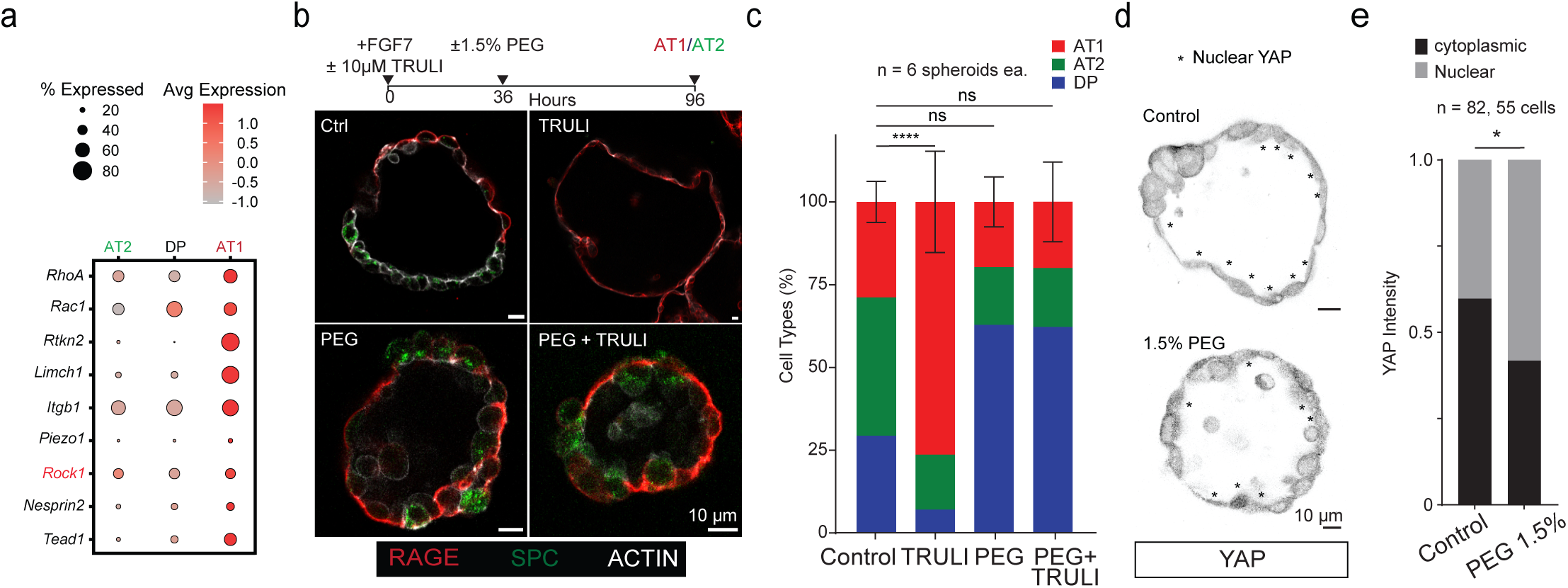
(A) scRNA seq analysis of genes in the alveolar epithelium associated with mechanotransduction and architectural remodeling. (B) Representative images of DPs cultured in control or TRULI (Lats inhibitor) media with or without PEG (scale bars, 10μm). (C) Quantification of AT1 cells from (B). Data presented as mean ± SD, p-values ****p < 0.0001 and ns (p ≥ 0.05) were calculated using one-way ANOVA. n = 6 spheroids for each condition. (D) Representative images of cytoplasmic to nuclear YAP intensity in PEG treated spheroids vs control (scale bars, 10μm). (E) Quantification of (D) nuclear and cytoplasmic YAP intensity (p-value = 0.038 by Chi Square Test). n = 83 and 55 cells for Control and 1.5% PEG conditions, respectively.

**Supplemental Figure 4:**
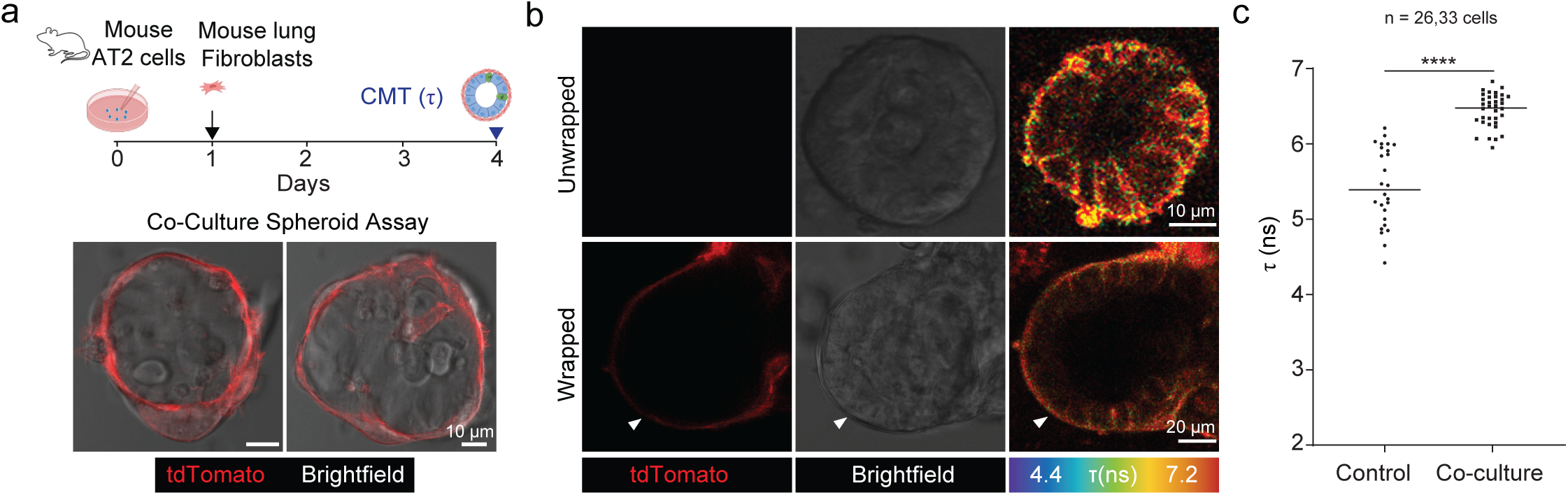
(A) Top: Schematic detailing the co-culturing mouse AT2 cells with labeled mouse fibroblasts. Bottom: Representative images of AT2 spheroids wrapped with labeled mouse fibroblasts. Scale bar = 15μm. (B) Representative Flipper-TR and Brightfield images of wrapped and unwrapped spheroids. Arrow denotes representative wrapped region. Scale bar = 15μm. (C) Graph showing quantification of Flipper-TR lifetime by τ-gating of wrapped vs unwrapped spheroids. Data represents the means ± SDs of 3 independent experiments. p value ****<0.0001 was calculated using a Student’s t-test. n = 26 and 35 for the unwrapped and wrapped groups, respectively.

**Supplemental Figure 5:**
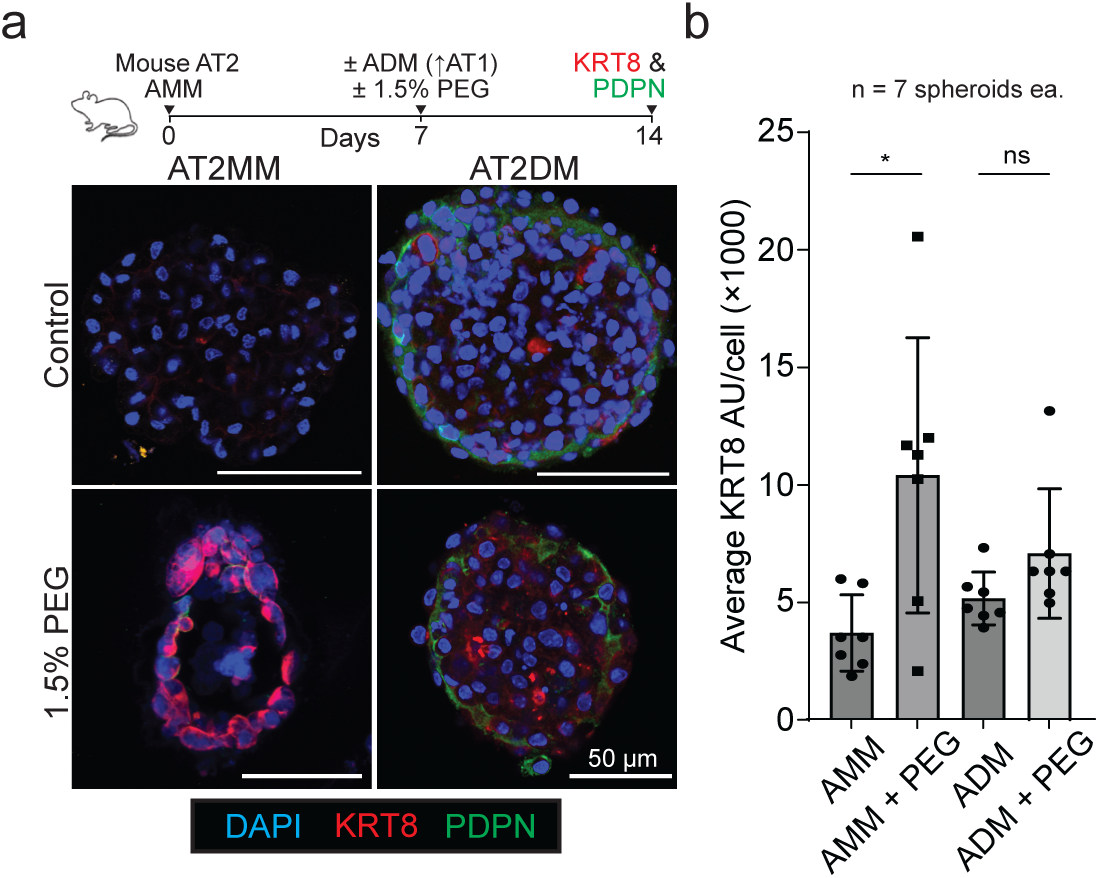
(A) Mouse AT2 organoids cultured in AMM or ADM ± 1.5% PEG and stained for KRT8 and PDPN. Scale bar, 50 µm. (B) Quantification of KRT8 intensity per cell across conditions (n = 7 spheroids each)

## Acknowledgements

We would like to extend special thanks to our students Thea Hartley, Jack Carlton, and Gamaliel Taengwa for their work on this project. This work was supported by the US National Institute of Health grants 5R00HL127267-04 and 1R01HL171056.

## Author contributions

GA, DG, and DGB conceived the study and designed the experiments. GA and DG performed the bulk of experiments. AM, MM, and IG aided in the collection and use of human fetal tissue samples; YH and MG performed the optical trap measurements; JC contributed the co-culture protocol; AM made and analyzed the data deriving from the iMC-linker virus; AO, IK, and EM provided operational support. GA, DG, and EG performed cell culture experiments, analyzed and interpreted the data. DGB, GA, and DG wrote the paper.

## Declaration of generative AI and AI-assisted technologies in the manuscript preparation process

During the preparation of this work the authors used ChatGPT in order to edit the manuscript for cogency and correctness. After using this tool/service, the authors reviewed and edited the content as needed and take full responsibility for the content of the published article.

## Data/Code Availability

All original microscopy and imaging data (including confocal .lif files, etc.) are available upon reasonable request. All original code for data processing, pathway inference, and figure generation will be made available on GitHub upon publication of this manuscript. The analysis was performed using R (v4.4.2) and the following key packages: Seurat (v5.1.0) and PROGENy (v1.28.0).

## References

Ali, G., Zhang, M., Zhao, R., Jain, K.G., Chang, J., Komatsu, S., Zhou, B., Liang, J., Matthay, M.A., and Ji, H.L. (2021). Fibrinolytic niche is required for alveolar type 2 cell-mediated alveologenesis via a uPA-A6-CD44(+)-ENaC signal cascade. Signal Transduct Target Ther 6, 97.

Bergert, M., Lembo, S., Sharma, S., Russo, L., Milovanovic, D., Gretarsson, K.H., Bormel, M., Neveu, P.A., Hackett, J.A., Petsalaki, E., et al. (2021). Cell Surface Mechanics Gate Embryonic Stem Cell Differentiation. Cell Stem Cell 28, 209–216 e204.

Bijonowski, B.M., Park, J., Bergert, M., Teubert, C., Diz-Muñoz, A., Galic, M., and Wegner, S.V. (2025). Intercellular adhesion boots collective cell migration through elevated membrane tension. Nature Communications 16, 1588.

Borghi, N., Sorokina, M., Shcherbakova, O.G., Weis, W.I., Pruitt, B.L., Nelson, W.J., and Dunn, A.R. (2012). E-cadherin is under constitutive actomyosin-generated tension that is increased at cell-cell contacts upon externally applied stretch. Proc Natl Acad Sci U S A 109, 12568–12573.

Brownfield, D.G., de Arce, A.D., Ghelfi, E., Gillich, A., Desai, T.J., and Krasnow, M.A. (2022). Alveolar cell fate selection and lifelong maintenance of AT2 cells by FGF signaling. Nat Commun 13, 7137.

Chan, C.J., Costanzo, M., Ruiz-Herrero, T., Monke, G., Petrie, R.J., Bergert, M., Diz-Munoz, A., Mahadevan, L., and Hiiragi, T. (2019). Hydraulic control of mammalian embryo size and cell fate. Nature 571, 112–116.

Choi, J., Park, J.E., Tsagkogeorga, G., Yanagita, M., Koo, B.K., Han, N., and Lee, J.H. (2020). Inflammatory Signals Induce AT2 Cell-Derived Damage-Associated Transient Progenitors that Mediate Alveolar Regeneration. Cell Stem Cell 27, 366–382 e367.

Colom, A., Derivery, E., Soleimanpour, S., Tomba, C., Molin, M.D., Sakai, N., Gonzalez-Gaitan, M., Matile, S., and Roux, A. (2018). A fluorescent membrane tension probe. Nat Chem 10, 1118–1125.

Da Silva André, G., and Labouesse, C. (2024). Mechanobiology of 3D cell confinement and extracellular crowding. Biophysical Reviews 16, 833–849.

De Belly, H., Stubb, A., Yanagida, A., Labouesse, C., Jones, P.H., Paluch, E.K., and Chalut, K.J. (2021). Membrane Tension Gates ERK-Mediated Regulation of Pluripotent Cell Fate. Cell Stem Cell 28, 273–284 e276.

Desai, R.A., Gao, L., Raghavan, S., Liu, W.F., and Chen, C.S. (2009). Cell polarity triggered by cell-cell adhesion via E-cadherin. J Cell Sci 122, 905–911.

Desai, T.J., Brownfield, D.G., and Krasnow, M.A. (2014). Alveolar progenitor and stem cells in lung development, renewal and cancer. Nature 507, 190–194.

Diz-Munoz, A., Fletcher, D.A., and Weiner, O.D. (2013). Use the force: membrane tension as an organizer of cell shape and motility. Trends Cell Biol 23, 47–53.

Elosegui-Artola, A., Andreu, I., Beedle, A.E.M., Lezamiz, A., Uroz, M., Kosmalska, A.J., Oria, R., Kechagia, J.Z., Rico-Lastres, P., Le Roux, A.L., et al. (2017). Force Triggers YAP Nuclear Entry by Regulating Transport across Nuclear Pores. Cell 171, 1397–1410 e1314.

Gillich, A., St. Julien, K.R., Brownfield, D.G., Travaglini, K.J., Metzger, R.J., and Krasnow, M.A. (2021). Alveoli form directly by budding led by a single epithelial cell. bioRxiv, 2021.2012.2025.474174.

Gokey, J.J., Snowball, J., Sridharan, A., Sudha, P., Kitzmiller, J.A., Xu, Y., and Whitsett, J.A. (2021). YAP regulates alveolar epithelial cell differentiation and AGER via NFIB/KLF5/NKX2-1. iScience 24, 102967.

Goodwin, K., Mao, S., Guyomar, T., Miller, E., Radisky, D.C., Kosmrlj, A., and Nelson, C.M. (2019). Smooth muscle differentiation shapes domain branches during mouse lung development. Development 146.

Han, Y.L., Ronceray, P., Xu, G., Malandrino, A., Kamm, R.D., Lenz, M., Broedersz, C.P., and Guo, M. (2018). Cell contraction induces long-ranged stress stiffening in the extracellular matrix. Proc Natl Acad Sci U S A 115, 4075–4080.

Jiang, P., Gil de Rubio, R., Hrycaj, S.M., Gurczynski, S.J., Riemondy, K.A., Moore, B.B., Omary, M.B., Ridge, K.M., and Zemans, R.L. (2020). Ineffectual Type 2-to-Type 1 Alveolar Epithelial Cell Differentiation in Idiopathic Pulmonary Fibrosis: Persistence of the KRT8(hi) Transitional State. Am J Respir Crit Care Med 201, 1443–1447.

Kobayashi, Y., Tata, A., Konkimalla, A., Katsura, H., Lee, R.F., Ou, J., Banovich, N.E., Kropski, J.A., and Tata, P.R. (2020). Persistence of a regeneration-associated, transitional alveolar epithelial cell state in pulmonary fibrosis. Nature Cell Biology 22, 934–946.

Konishi, S., Tata, A., and Tata, P.R. (2022). Defined conditions for long-term expansion of murine and human alveolar epithelial stem cells in three-dimensional cultures. STAR Protoc 3, 101447.

Li, J., Wang, Z., Chu, Q., Jiang, K., Li, J., and Tang, N. (2018). The Strength of Mechanical Forces Determines the Differentiation of Alveolar Epithelial Cells. Developmental Cell 44, 297–312.e295.

Li, Y., Chen, M., Hu, J., Sheng, R., Lin, Q., He, X., and Guo, M. (2021). Volumetric Compression Induces Intracellular Crowding to Control Intestinal Organoid Growth via Wnt/beta-Catenin Signaling. Cell Stem Cell 28, 63–78 e67.

Liberti, D.C., Kremp, M.M., Liberti, W.A., 3rd, Penkala, I.J., Li, S., Zhou, S., and Morrisey, E.E. (2021). Alveolar epithelial cell fate is maintained in a spatially restricted manner to promote lung regeneration after acute injury. Cell Rep 35, 109092.

Lim, K., Donovan, A.P.A., Tang, W., Sun, D., He, P., Pett, J.P., Teichmann, S.A., Marioni, J.C., Meyer, K.B., Brand, A.H., et al. (2023). Organoid modeling of human fetal lung alveolar development reveals mechanisms of cell fate patterning and neonatal respiratory disease. Cell Stem Cell 30, 20–37 e29.

Lim, K., and Rawlins, E.L. (2024). Protocol for the derivation and alveolar type 2 differentiation of late-stage lung tip progenitors from the developing human lungs. STAR Protoc 5, 103201.

Lopez, C.A., de Vries, A.H., and Marrink, S.J. (2011). Molecular mechanism of cyclodextrin mediated cholesterol extraction. PLoS Comput Biol 7, e1002020.

Nabhan, A.N., Brownfield, D.G., Harbury, P.B., Krasnow, M.A., and Desai, T.J. (2018). Single-cell Wnt signaling niches maintain stemness of alveolar type 2 cells. Science 359, 1118–1123.

Nantie, L.B., Young, R.E., Paltzer, W.G., Zhang, Y., Johnson, R.L., Verheyden, J.M., and Sun, X. (2018). Lats1/2 inactivation reveals Hippo function in alveolar type I cell differentiation during lung transition to air breathing. Development 145.

Pontes, B., Monzo, P., and Gauthier, N.C. (2017). Membrane tension: A challenging but universal physical parameter in cell biology. Semin Cell Dev Biol 71, 30–41.

Riggi, M., Bourgoint, C., Macchione, M., Matile, S., Loewith, R., and Roux, A. (2019). TORC2 controls endocytosis through plasma membrane tension. J Cell Biol 218, 2265–2276.

Rouven Bruckner, B., Pietuch, A., Nehls, S., Rother, J., and Janshoff, A. (2015). Ezrin is a Major Regulator of Membrane Tension in Epithelial Cells. Sci Rep 5, 14700.

Sawhney, A.S., Deskin, B.J., Cai, J., Gibbard, D., Ali, G., Utoft, A., Qi, X., Olson, A., Hausman, H., Sabol, L., et al. (2025). A molecular circuit regulates fate plasticity in emerging and adult AT2 cells. Nat Commun 16, 8924.

Schindelin, J., Arganda-Carreras, I., Frise, E., Kaynig, V., Longair, M., Pietzsch, T., Preibisch, S., Rueden, C., Saalfeld, S., Schmid, B., et al. (2012). Fiji: an open-source platform for biological-image analysis. Nat Methods 9, 676–682.

Shiraishi, K., Shah, P.P., Morley, M.P., Loebel, C., Santini, G.T., Katzen, J., Basil, M.C., Lin, S.M., Planer, J.D., Cantu, E., et al. (2023). Biophysical forces mediated by respiration maintain lung alveolar epithelial cell fate. Cell 186, 1478–1492 e1415.

Sorkin, A., and von Zastrow, M. (2009). Endocytosis and signalling: intertwining molecular networks. Nat Rev Mol Cell Biol 10, 609–622.

Strunz, M., Simon, L.M., Ansari, M., Kathiriya, J.J., Angelidis, I., Mayr, C.H., Tsidiridis, G., Lange, M., Mattner, L.F., Yee, M., et al. (2020). Alveolar regeneration through a Krt8+ transitional stem cell state that persists in human lung fibrosis. Nat Commun 11, 3559.

Sun, D., Llora Batlle, O., van den Ameele, J., Thomas, J.C., He, P., Lim, K., Tang, W., Xu, C., Meyer, K.B., Teichmann, S.A., et al. (2022). SOX9 maintains human foetal lung tip progenitor state by enhancing WNT and RTK signalling. EMBO J 41, e111338.

Treutlein, B., Brownfield, D.G., Wu, A.R., Neff, N.F., Mantalas, G.L., Espinoza, F.H., Desai, T.J., Krasnow, M.A., and Quake, S.R. (2014). Reconstructing lineage hierarchies of the distal lung epithelium using single-cell RNA-seq. Nature 509, 371–375.

Valenta, T., Hausmann, G., and Basler, K. (2012). The many faces and functions of β-catenin. Embo j 31, 2714–2736.

Wang, F., Ting, C., Riemondy, K.A., Douglas, M., Foster, K., Patel, N., Kaku, N., Linsalata, A., Nemzek, J., Varisco, B.M., et al. (2023). Regulation of epithelial transitional states in murine and human pulmonary fibrosis. J Clin Invest 133.

Wang, Y., Wang, L., Ma, S., Cheng, L., and Yu, G. (2024). Repair and regeneration of the alveolar epithelium in lung injury. FASEB J 38, e23612.

Watson, J., Ferguson, H.R., Brady, R.M., Ferguson, J., Fullwood, P., Mo, H., Bexley, K.H., Knight, D., Howell, G., Schwartz, J.M., et al. (2022). Spatially resolved phosphoproteomics reveals fibroblast growth factor receptor recycling-driven regulation of autophagy and survival. Nat Commun 13, 6589.

Wu, X.S., Elias, S., Liu, H., Heureaux, J., Wen, P.J., Liu, A.P., Kozlov, M.M., and Wu, L.G. (2017). Membrane Tension Inhibits Rapid and Slow Endocytosis in Secretory Cells. Biophys J 113, 2406–2414.

Young, R.E., Jones, M.-K., Hines, E.A., Li, R., Luo, Y., Shi, W., Verheyden, J.M., and Sun, X. (2020a). Smooth Muscle Differentiation Is Essential for Airway Size, Tracheal Cartilage Segmentation, but Dispensable for Epithelial Branching. Developmental Cell 53, 73–85.e75.

Zepp, J.A., Morley, M.P., Loebel, C., Kremp, M.M., Chaudhry, F.N., Basil, M.C., Leach, J.P., Liberti, D.C., Niethamer, T.K., Ying, Y., et al. (2021). Genomic, epigenomic, and biophysical cues controlling the emergence of the lung alveolus. Science 371.

Zepp, J.A., Zacharias, W.J., Frank, D.B., Cavanaugh, C.A., Zhou, S., Morley, M.P., and Morrisey, E.E. (2017). Distinct Mesenchymal Lineages and Niches Promote Epithelial Self-Renewal and Myofibrogenesis in the Lung. Cell 170, 1134–1148.e1110.

Zhang, Y., Angiulli, G., Martinac, B., Cox, C.D., and Walz, T. (2021). Cyclodextrins for structural and functional studies of mechanosensitive channels. J Struct Biol X 5, 100053.

